# Fluorescence polarization-based fragment screen identifies inhibitors of APOBEC3A and APOBEC3B

**DOI:** 10.64898/2026.07.31.742089

**Authors:** Julianna S. Brunello, Hannah R. Allegakoen, Weicheng Li, Jezrael L. Revalde, Rohit Bhadoria, Colton M. Sanders, Michelle R. Arkin, Jonathan M. L. Ostrem

**Author notes:** Corresponding Authors: Jonathan M. L. Ostrem – Department of Medicine, University of California, San Francisco, San Francisco, California 94143.,; Hannah R. Allegakoen – Department of Pharmaceutical Chemistry, University of California, San Francisco, San Francisco, California 94143.

## Abstract

APOBEC3A and APOBEC3B are antiviral cytidine deaminases found to drive cancer-associated mutagenesis, contributing to tumor evolution and therapeutic resistance across multiple cancer types. Inhibiting these enzymes holds promise for prolonging response to a wide range of cancer therapies by delaying development of resistance. However, APOBEC3A and APOBEC3B remain challenging drug targets, with no potent and selective small molecule inhibitors reported. Here, we use a fluorescence polarization-based assay to identify small molecules inhibitors of the APOBEC3A-single-stranded DNA interaction. From a library of 2,400 disulfide compounds, we identified 64 hits (mean polarization +/- 3 sigma, hit rate of 2.7%). Intact protein mass spectrometry revealed that a subset of compounds covalently engages A3A at cysteine 64, including Compounds 1 and 2. Compounds 1 and 2 disrupt APOBEC3A/APOBEC3B-single-stranded DNA interactions and inhibit APOBEC3A/APOBEC3B deaminase activity in a dose-dependent manner, with micromolar IC_50_. Surprisingly, inhibition of APOBEC3A/APOBEC3B by Compounds 1 and 2 is independent of covalent tethering to cysteine, suggesting a predominantly non-covalent mode of binding. Together, these studies establish an integrated workflow for APOBEC ligand discovery and identify Compounds 1 and 2 as starting points for developing chemical probes to investigate APOBEC-driven mutagenesis and therapeutic resistance.

---

Genomic instability is a defining feature of cancer that facilitates tumor adaptation and treatment resistance. Apolipoprotein B mRNA-editing catalytic subunit-like 3 (APOBEC3) cytidine deaminases are central drivers of mutagenesis and genome instability across a wide variety of human cancers.^1^ By catalyzing the deamination of cytosine to uracil, APOBEC3 enzymes generate C>T/G mutations at 5′-TC-3′ motifs.^2,3^ APOBEC signatures are among the most prevalent mutational signatures in cancer genomes, second only to those associated with aging.^4^

APOBEC3 activity has been linked to tumor evolution and acquired therapeutic resistance across multiple cancers, primarily driven by two homologous isoforms, APOBEC3A (A3A) and APOBEC3B (A3B).^5–7^ In non-small cell lung cancer (NSCLC), treatment with tyrosine kinase inhibitors (TKIs) has been associated with increased APOBEC mutagenesis in patient tumors, suggesting that therapeutic pressure can amplify this mutational process. Consistent with this model, genetic depletion of A3A and A3B impairs the ability of NSCLC cell lines to acquire drug-tolerant persister states or develop resistance to targeted therapy.^5,8,9^ In breast cancer, elevated A3A/A3B expression or APOBEC mutational activity has similarly been associated with more aggressive disease and worse clinical outcomes.^10–13^ Together, these findings supporting a model in which A3A and A3B accelerate tumor evolution and development of therapeutic resistance.

A3A and A3B are nonessential and often overexpressed in tumors, making them attractive targets to delay acquired resistance.^1^^4,15^ Chemical probes that modulate A3A and/or A3B activity would enable direct investigation of their role in acquired resistance and evaluation of APOBEC inhibition as a viable strategy to delay tumor adaptation.

Efforts to develop APOBEC inhibitors have employed both substrate-mimetic and small-molecule strategies. Previous studies focused on single-stranded DNA substrates containing cytidine analogs such as zebularine, which act as transition-state mimics and inhibit APOBEC-mediated deamination.^16–21^ While these compounds established proof-of-concept for APOBEC inhibition, their utility is limited by modest potency, poor cellular permeability, and inherent challenges associated with oligonucleotide-based therapeutics.^22^ More recently, small-molecule approaches have explored covalent, allosteric, and structure-based inhibition. The first reported small-molecule APOBEC inhibitor targeted APOBEC3G through covalent engagement of a cysteine residue proximal to the catalytic site, demonstrating that APOBEC enzymes are chemically tractable targets.^23^ Subsequent studies identified compounds targeting putative allosteric sites in A3B and used large-scale virtual screening to discover inhibitors of AID, A3A, and A3B.^24,25^ Despite these advances, potent and selective chemical probes for A3A and A3B remain limited. This unmet need motivates the development of new screening strategies to identify ligands capable of modulating APOBEC activity and interrogating the role of APOBEC-mediated mutagenesis in cancer.

Here, we report discovery of small molecule inhibitors of APOBEC3 family cytidine deaminases from a disulfide tethering library. Tethering screens are typically run using mass spectrometry to identify fragments that covalently bind a target through disulfide bond formation. Here we instead elected to pursue a functional screen using fluorescence polarization to focus specifically on identifying fragments that altered DNA binding. Surprisingly, while a subset of hit fragments tether to cysteine-64 in A3A, we find that covalent interaction is not required for inhibition of DNA binding. Encouragingly, two hit fragments, compound 1 and compound 2 inhibit deaminase activity of purified A3A and lysates expressing A3A, A3B, A3F or A3G. Molecular docking suggests a mode of non-covalent binding in which the dichlorophenol functional group coordinates the active size zinc. However, neither compound inhibits histone deacetylases, another class of metalloenzyme containing a catalytic zinc ion, suggesting compounds 1 and 2 are not non-specific inhibitors of zinc-dependent metalloenzymes. Our studies provide promising starting points for APOBEC probe development and drug discovery efforts.

## RESULTS

### High-Throughput Fluorescence Polarization Screening Identifies Modulators of A3A- ssDNA Binding

Despite the central role of APOBEC3A (A3A) and APOBEC3B (A3B) in tumor evolution and therapeutic resistance, there are no approved inhibitors and few chemical tools targeting these enzymes.^26,27^ Building on our previous fragment-based discovery of KRAS ligands, we applied cysteine tethering to identify ligands targeting A3A and A3B.^28^ A3A and A3B contain surface-exposed cysteine residues near the ssDNA-binding interface (C64 and C247, respectively) that are distinct from the catalytic zinc-binding site (Figure 1a).^29^ Recent *in silico* studies identified covalent ligands targeting A3B C247, demonstrating the accessibility of this residue.^24^ Because C247 is analogous to C64 in A3A, we hypothesized that covalent targeting of C64 could disrupt A3A–ssDNA binding.^30^

**Figure 1.**
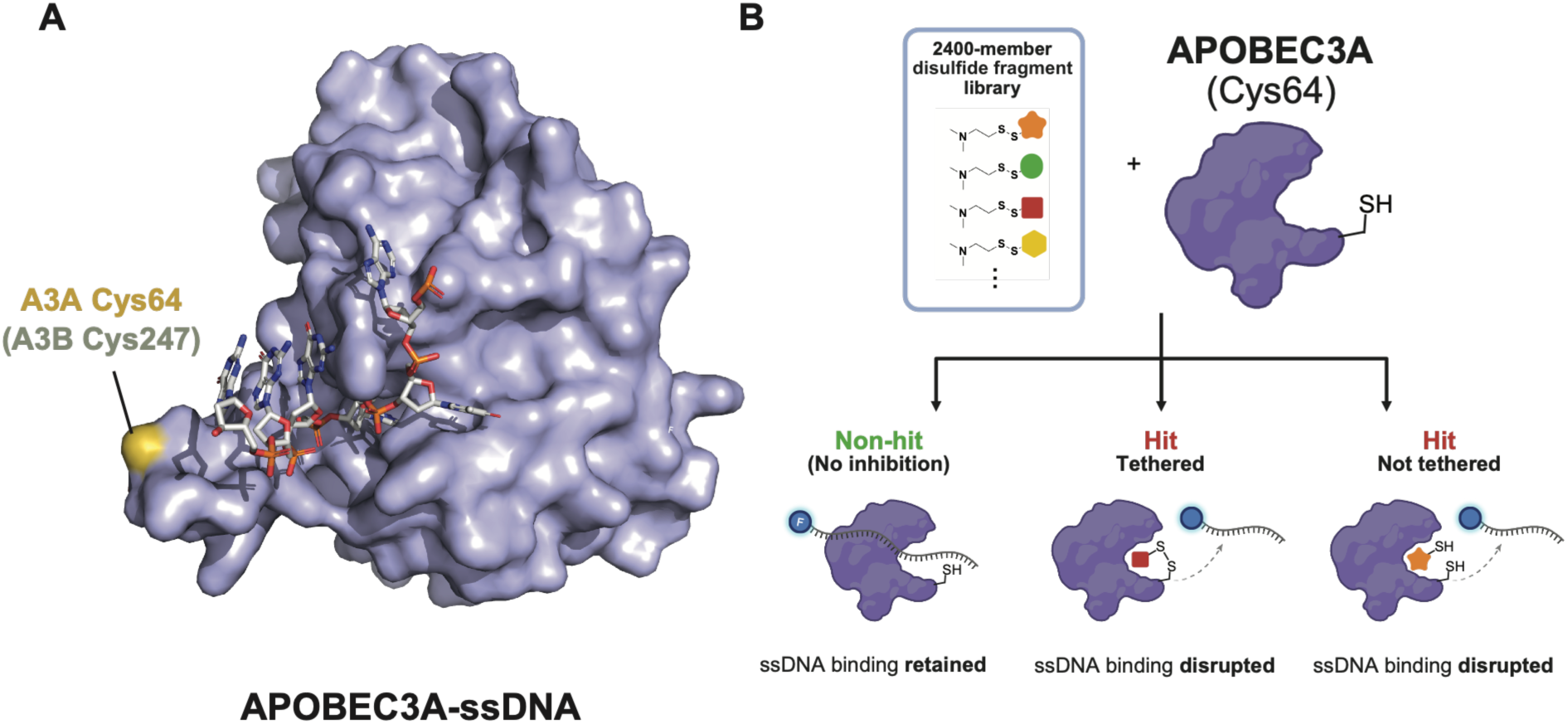
Strategy for identification of A3A inhibitors (A) APOBEC3A*^E72A^*-ssDNA structure (PDB: 5SWW) highlighting Cys64 (gold) and the analogous APOBEC3B residue Cys247 (gray). (B) Schematic of the fluorescence polarization–based disulfide fragment screen. 2,400-member disulfide fragment library was screened for compounds that disrupt A3A–ssDNA binding. Binding of A3A to fluorescently labeled ssDNA increases polarization, whereas compounds that disrupt the complex reduce polarization and are identified as hits.

Traditional disulfide tethering screens identify hits by mass spectrometry without requiring them to have functional effects. To focus our search on functional compounds that disrupt the interaction between A3A/A3B and ssDNA, we developed a fluorescence polarization (FP) assay based on validated methods for measuring APOBEC binding to ssDNA.^17,31–33^ Binding of fluorescently labeled ssDNA to A3A produces an increase in polarization due to reduced rotational diffusion of the bound oligonucleotide, whereas compounds that disrupt A3A– ssDNA complex formation decrease the polarization signal (Figure 1b). To eliminate turnover and allow stable complex formation, we performed initially screening against recombinant human A3A^E72A^, a catalytically inactive mutant.^34,35^ The protein was expressed in *E. coli* as N-terminal His_6_-GST fusion, with removal of both tags prior to final purification.

Optimization of oligonucleotide sequence and pH and selection of Cy5 dye to minimize of fluorescence interference, yielded a robust FP assay for high-throughput screening at physiologic pH 7.5 (Figure S1A-B). As expected, under these conditions, A3A^E72A^ showed preferential binding to TC-containing substrate (Kd = 326 nM) relative to TU- or TA-containing controls (Kd = 750 nM and 350 mM, respectively) (Figure S1C).^31,32^ For screening purposes under these optimized conditions we selected the EC_90_ concentration of 3.2 μM A3A^E72A^ (Figure 2a), and assay robustness was confirmed with a Z’ factor greater than 0.7.

**Figure 2.**
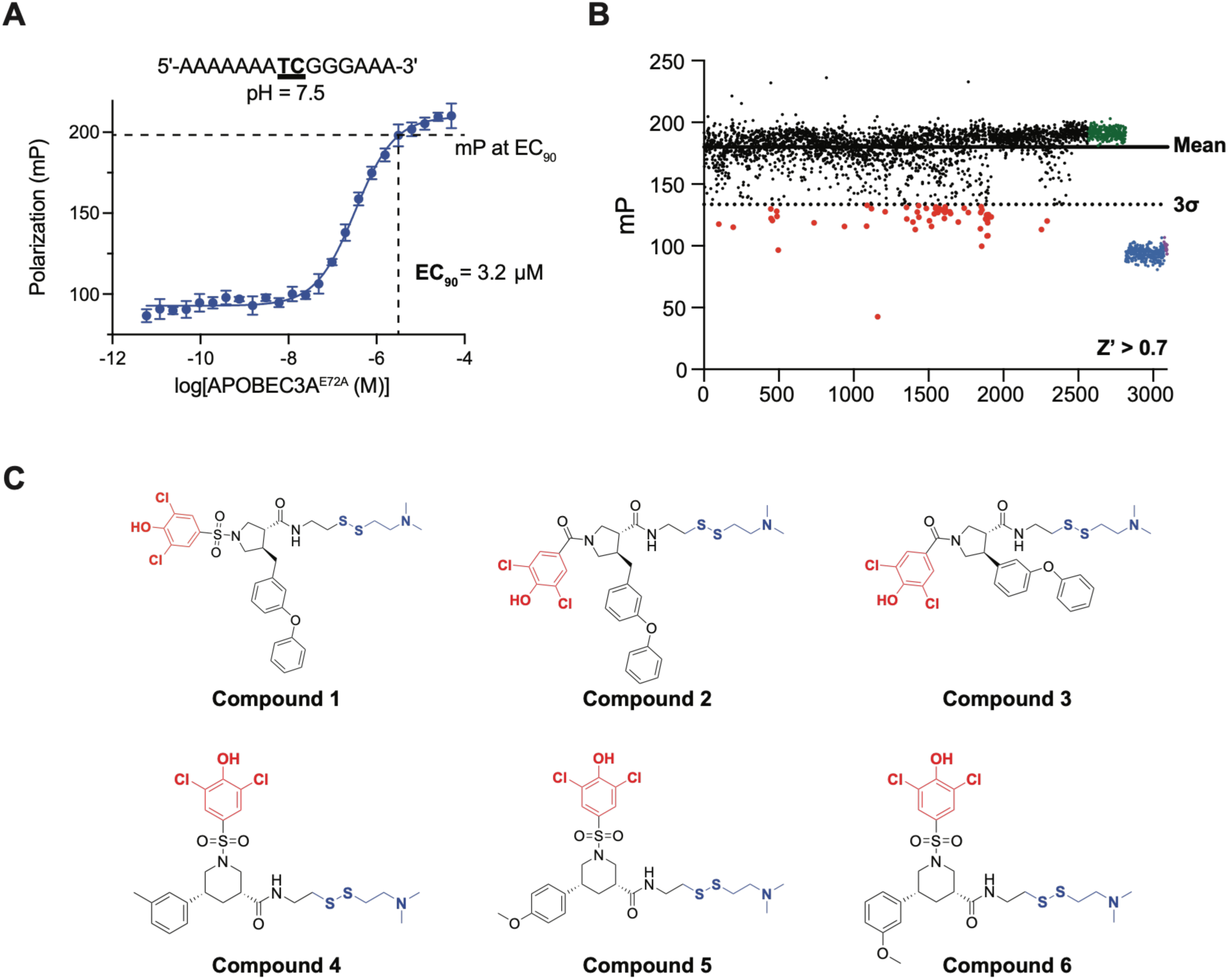
High-throughput fluorescence polarization screening identifies dichlorophenol-containing A3A inhibitors. (A) Binding of A3A^E72A^ to a fluorescently labeled ssDNA substrate under optimized assay conditions (pH 7.5). The EC₉₀ concentration (3.2 μM) was selected for screening. (B) Primary screening results for the 2,400-member disulfide fragment library. Hits (red) were defined as compounds producing polarization values >3 standard deviations below the mean. Free tracer (blue), A3A^E72A^–ssDNA complex (green), and MN1 (purple) are shown as controls. (C) Representative dichlorophenol-containing disulfide fragments (DCPs) identified in the primary screen.

To identify disulfide fragments that block the interaction between A3A^E72A^ and ssDNA, we screened a 2,400-member disulfide fragment library (Figure 2b). MN1, a broad inhibitor of nucleic acid-binding proteins, was included as a pharmacological control showing reduction of mP in the range of tracer oligonucleotide in the absence of protein (Figure 2b).^36^

By defining hits as fragments producing a change in mP greater than three standard deviations from the mean, we identified 64 hit fragments, corresponding to a hit rate of ∼2.5% (Figure 2b). Analysis of hit structures revealed an enrichment of dichlorophenol (DCP)- containing fragments, with 26 of 60 DCP-containing compounds in the library identified as hits (Figure 2c).

### A subset of dichlorophenol fragments covalently tether to A3A C64

A3A contains two solvent-exposed cysteines, C64 and C171 (Figure 3a). Intact-protein LC– MS following 1mM iodoacetamide treatment showed an ∼115 Da mass increase, consistent with alkylation of both cysteines and confirming that each residue is accessible for covalent modification (Figure 3a). Having established reactivity of C64, we evaluated all 64 disulfide fragment hits for covalent modification of A3A by LC-MS at 1 μM A3A^E72A^, 200 μM fragment, and 0.5 mM tris(2-carboxyethyl)phosphine (TCEP). Formation of a disulfide bond between a fragment and a cysteine residue on A3A is expected to produce a corresponding protein mass shift detectable by intact-protein LC-MS. Of the 64 FP hits, 10 DCP-containing fragments exhibited detectable covalent modification of A3A. Among these, compounds 1 and 2 produced the highest levels of labeling, reaching 48.7 ± 1.7% and 58.2 ± 3.2% covalent modification, respectively (mean ± SD) (Figure 3b).

**Figure 3.**
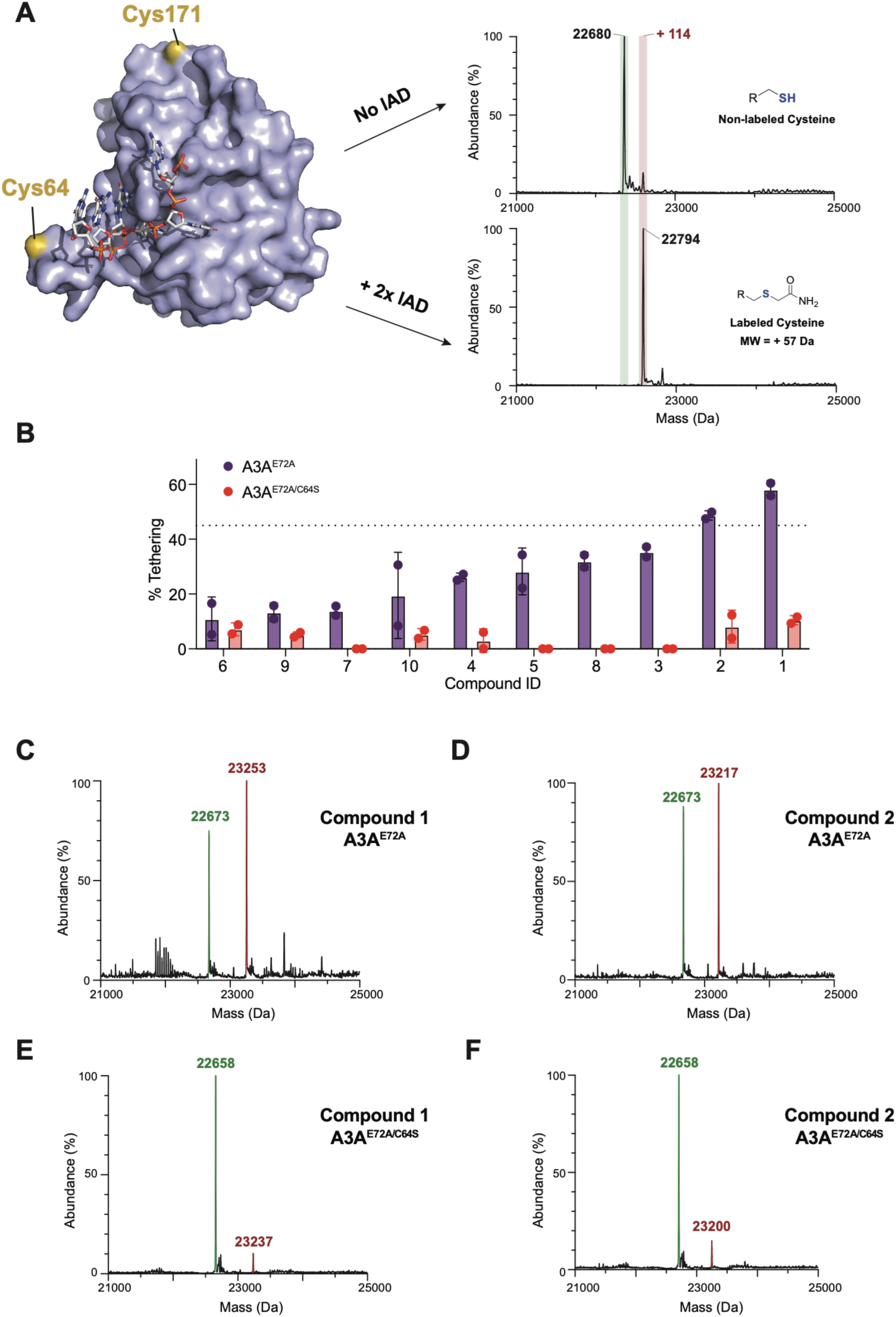
Identification of dichlorophenol-containing tethering hits that engage A3A through C64. (A) Structure of A3A highlighting the solvent-exposed cysteines C64 and C171 and LC–MS analysis following iodoacetamide (IAM) treatment showing alkylation of both residues (PDB: 5SWW). (B) Covalent labeling of A3A by the 10 dichlorophenol-containing fragments identified in the tethering screen. (C,D) LC–MS analysis of wild-type A3A incubated with Compounds 1 and 2, respectively, showing formation of protein–fragment adducts. (E,F) LC– MS analysis of A3A^E72A/C64S^ incubated with Compounds 1 and 2. Loss of the protein–fragment adduct demonstrates C64-dependent labeling.

Intact protein LC-MS analysis of A3A incubated with compounds 1 and 2 revealed additional mass peaks at +580 Da and +544 Da, respectively, indicating covalent tethering of a single fragment molecule per protein despite the presence of two accessible cysteine residues (Figure 3 c,d). These additional peaks were significantly less prominent when these compounds were instead incubated with the A3A^E72A/C64S^ mutant, suggesting modification occurs primarily at C64 (Figure 3 e,f).

### Compounds 1 and 2 inhibit APOBEC3A deaminase activity independent of C64 tethering

Based on their apparent interference with ssDNA binding, we wondered whether compounds 1 and 2 would block the deaminase activity of wild type A3A (A3A^WT^) or the tethering-impaired mutant A3A^C64S^. We expressed and purified recombinant A3A^WT^ and A3AC^64S^ from mammalian suspension cells. We then used an established gel-based cytosine deaminase assay to evaluate the effect of compounds 1 and 2 on A3A catalytic activity (Figure 4a).^37^ In this assay, fluorescently labeled ssDNA substrate is incubated with APOBEC enzyme and uracil DNA glycosylase (UDG). Upon deamination of cytosine to uracil by APOBEC, UDG excises the uracil base, enabling alkaline cleavage at the resulting abasic site to yield a shorter DNA fragment that can be resolved by gel electrophoresis.^23,37,38^ Inhibition of A3A catalytic activity results in reduced substrate cleavage.

**Figure 4.**
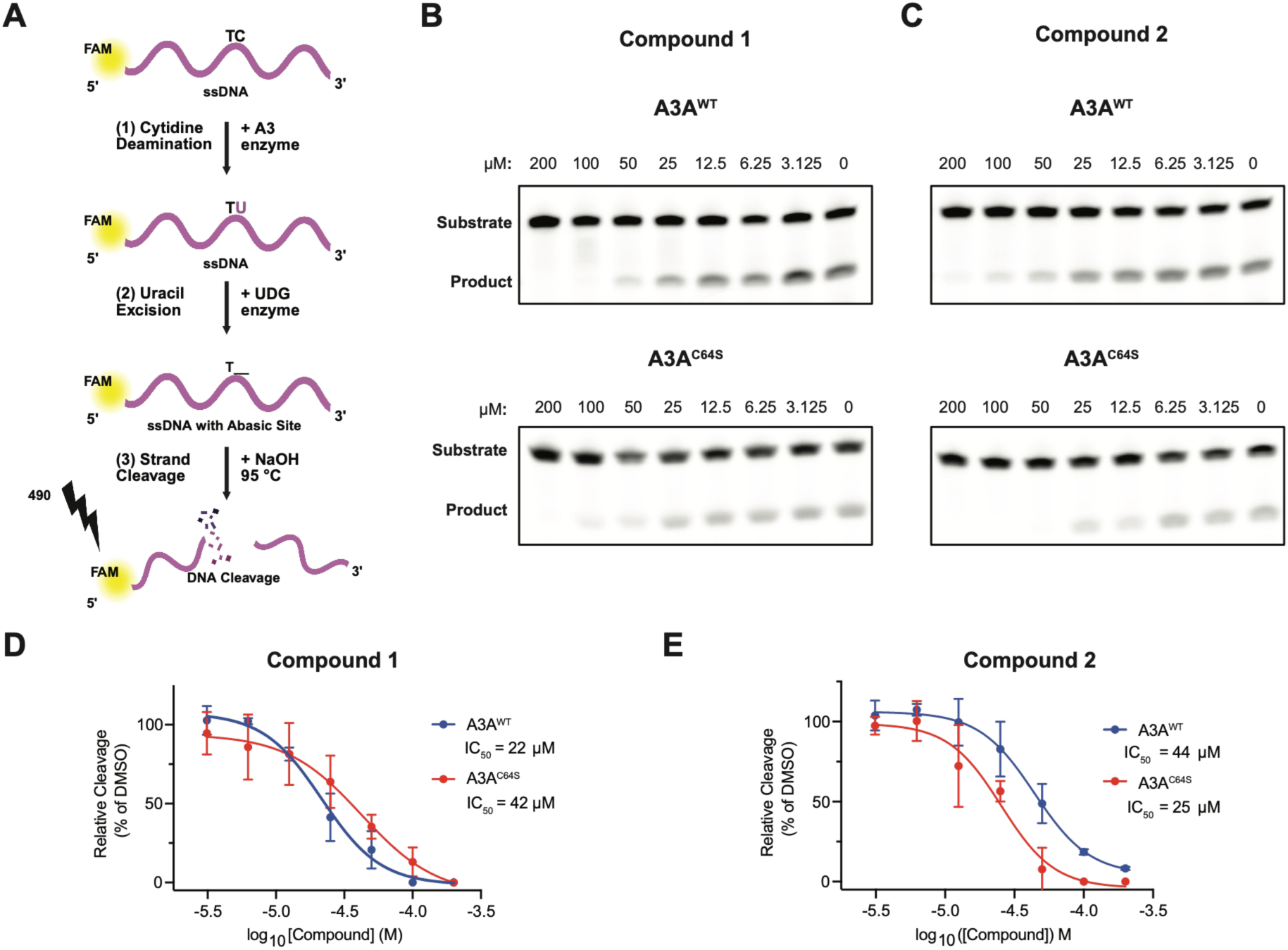
Lead DCP Compounds Inhibit A3A Deaminase Activity Independent of C64 Tethering (A) Schematic of the gel-based DNA deaminase assay. A3A deaminates a fluorescent ssDNA substrate, which is subsequently cleaved following UDG excision and alkaline treatment. (B) Dose-dependent inhibition of A3A^WT^ and A3A^C64S^ by Compound 1. Representative gels and dose-response curves are shown. IC₅₀ values were 22 μM and 42 μM for A3A^WT^ and A3A^C64S^, respectively. (C) Dose-dependent inhibition of A3A^WT^ and A3A^C64S^ by Compound 2. Representative gels and dose-response curves are shown. IC₅₀ values were 44 μM and 25 μM for A3A^WT^ and A3A^C64S^, respectively. (D,E) Quantification of the dose-response experiments shown in (B) and (C), respectively.

Compounds 1 and 2 both inhibited A3A^WT^ in a concentration-dependent manner, with compound 1 exhibiting slightly greater potency than compound 2 (apparent IC_50_ of 22 µM versus 44 µM) (Figure 4b,c). Reaction conditions, including A3A concentration, were optimized to achieve approximately 50% substrate conversion over the experimental timeframe. Additional assay optimization and validation experiments are shown in Supplementary Figures S2-S5. To determine whether enzymatic inhibition depended on covalent modification of C64, we repeated this assay using the A3A^C64S^ mutant. Surprisingly, both compounds retained inhibitory activity, with nearly identical apparent IC_50_ values compared with A3A^WT^ (Figure 4b, c). Compound 1 exhibited IC_50_ of 22 µM and 42 µM against A3A^WT^ and A3A^C64S^, respectively, whereas compound 2 exhibited IC_50_ of 44 µM and 25 µM against these respective enzymes (Figure 4 d,e). Under similar assay conditions, MN1 inhibited A3A^WT^ with an apparent IC_50_ of 0.53 µM (Figure S5). These results indicate that covalent tethering to C64 is dispensable for inhibition A3A catalytic activity and suggest that non- covalent interactions contribute substantially to the observed activity.

### Compounds 1 and 2 inhibit APOBEC3A and APOBEC3B deaminase activity in lysates

Like A3A, APOBEC3B (A3B) is a key source of genomic instability across multiple cancers.^10,41,5,40^ Full-length A3B has proven difficult to purify due to challenges associated with its N-terminal domain (NTD), and previous studies have demonstrated that the NTD contributes to catalytic efficiency and deamination activity.^44–46^ Therefore, to determine whether compounds 1 and 2 also inhibit A3B, we performed a modified version of our cytidine deaminase assay using lysates from HEK293T cells transiently overexpressing A3B. HEK293T cells exhibit negligible endogenous APOBEC3 expression, and activity in lysates can be attributed to the exogenously expression APOBEC enzyme (Figure S6).^37^ Assay conditions were again optimized by varying lysate input and selecting conditions that produced partial substrate turnover within the dynamic range of the assay (Figure S7).

Both compounds inhibited A3B deaminase activity in a concentration-dependent manner (Figure 5 a,b) with apparent IC₅₀ of 43 μM and 41 μM for compound 1 and compound 2, respectively (Figure 5 c,d). Similar to A3A, mutation of the cysteine analogous to C64 in A3A (A3B^C247S^) had little effect on inhibitory activity of either compound with only minor differences in potency observed for each compound against A3B^C247S^ versus A3B^WT^. These results are again consistent with cysteine tethering being dispensable for inhibition of APOBEC by compounds 1 and 2.

**Figure 5.**
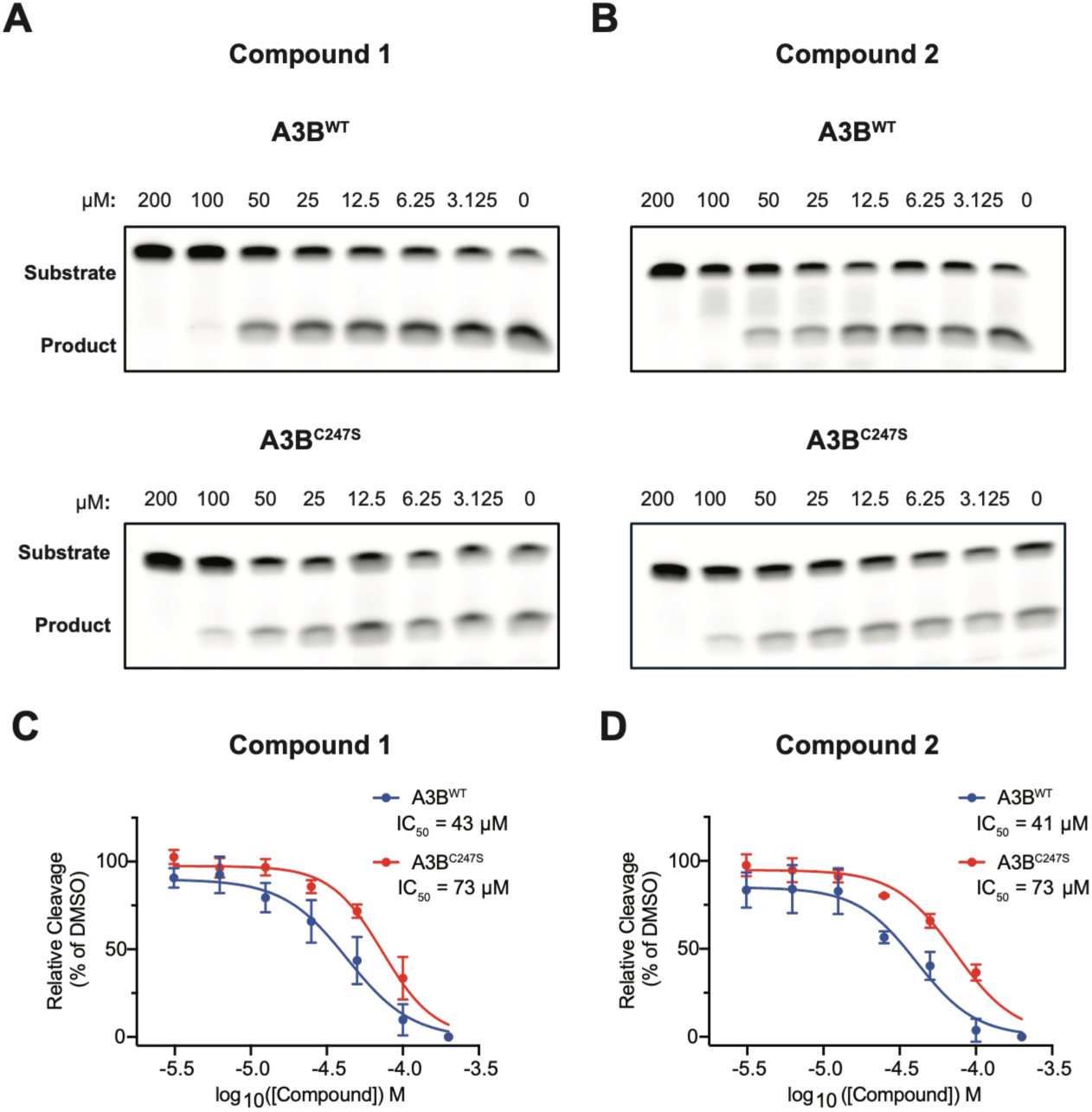
Compounds 1 and 2 Inhibit APOBEC3B Independent of C247 Tethering (A,B) Representative dose-response gels showing inhibition of A3B^WT^ and A3B^C247S^ by Compounds 1 and 2, respectively. (C) Quantification of Compound 1 inhibition in A3B^WT^ and A3B^C247S^. Apparent IC₅₀ values were 43 μM and 73 μM, respectively. (D) Quantification of Compound 2 inhibition in A3B^WT^ and A3B^C247S^. Apparent IC₅₀ values were 41 μM and 73 μM, respectively.

### Molecular docking suggests DCPs bind through coordination of active site zinc

To better understand the mode of binding of compounds 1 and 2, we performed docking studies using the Molecular Operating Environment (MOE). Docking of compound 1 to the structure of A3A^E72A^:ssDNA (PDB: 5SWW) predicted a consistent binding mode demonstrating coordination of the active site zinc ion by the dichlorophenol functional group, with docking scores ranging from S=-13.58 to -12.38 kcal/mol across the top five poses (Figure 6).

**Figure 6.**
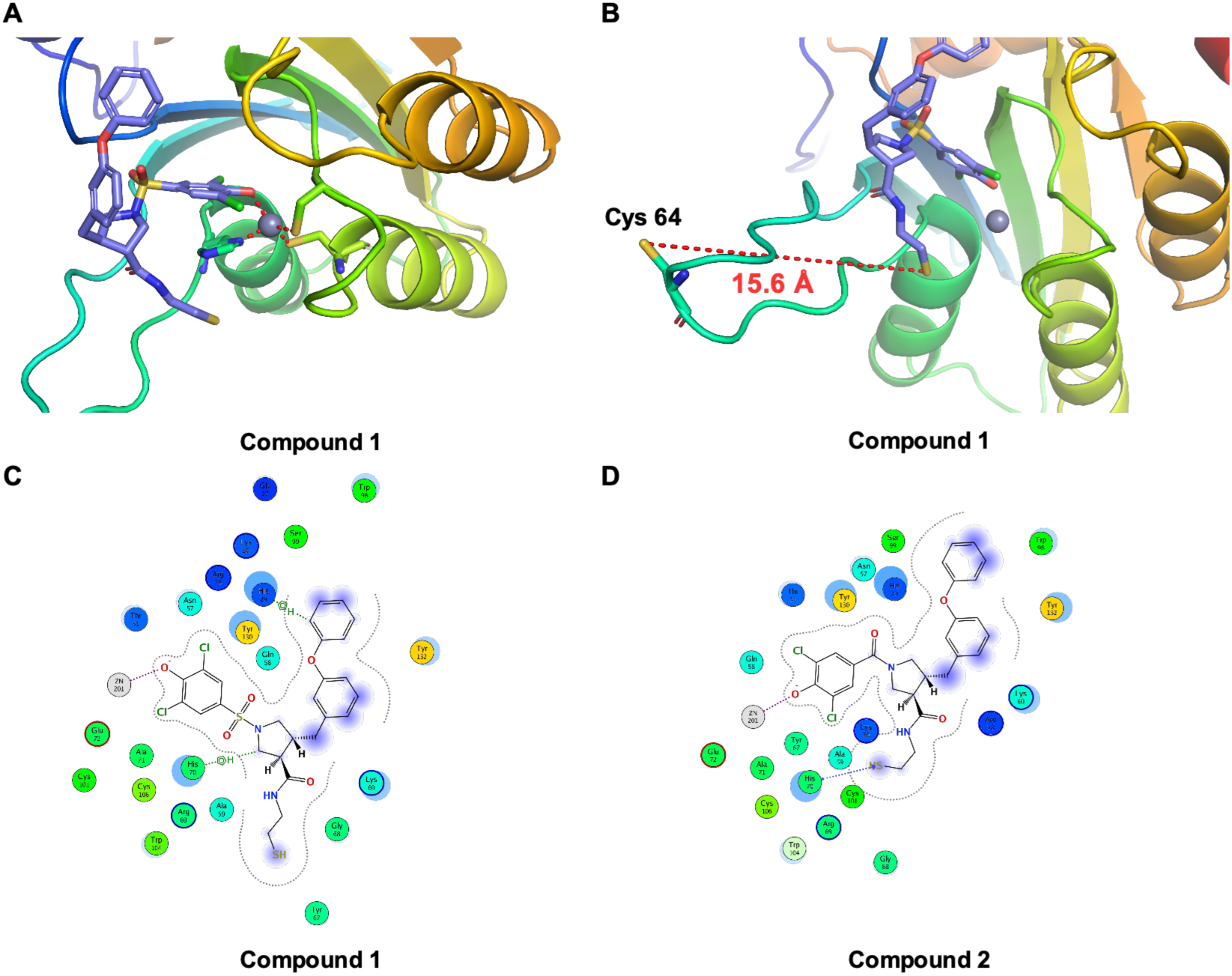
Docking Predicts Noncovalent Binding of Compounds 1 and 2 (A,B) Docking models of Compounds 1 and 2 in the A3A active site position the dichlorophenol group adjacent to the catalytic Zn²⁺. The disulfide is separated from Cys64 (∼15.6 Å), suggesting inhibition occurs through noncovalent binding rather than tethering. (C,D) Predicted ligand interactions for Compounds 1 and 2.

### Compounds 1 and 2 exhibit activity across multiple APOBEC family members

We next evaluated Compounds 1 and 2 against additional APOBEC3 (A3) family members. Although APOBEC3 enzymes share a conserved zinc-dependent cytidine deaminase fold, they differ in domain organization (Figure 7A).^47,48^ We evaluated A3A, A3B, A3G, and A3F using lysates from HEK293T cells expressing each enzyme. For A3A, apparent potency was reduced relative to purified protein, likely reflecting the increased complexity of the lysate environment. Both compounds inhibited A3A, A3B, A3G, and A3F, demonstrating broad activity across representative APOBEC3 family members (Figure 7B,C). To assess selectivity, Compounds 1 and 2 were counter-screened against Class I/II histone deacetylases (HDACs). Neither compound inhibited HDAC activity, whereas the positive control trichostatin A produced the expected inhibition (Figure S12). These findings indicate that the dichlorophenol scaffold does not act as a nonspecific zinc-metalloenzyme inhibitor and provides a promising starting point for medicinal chemistry to improve potency and engineer isoform selectivity.

**Figure 7.**
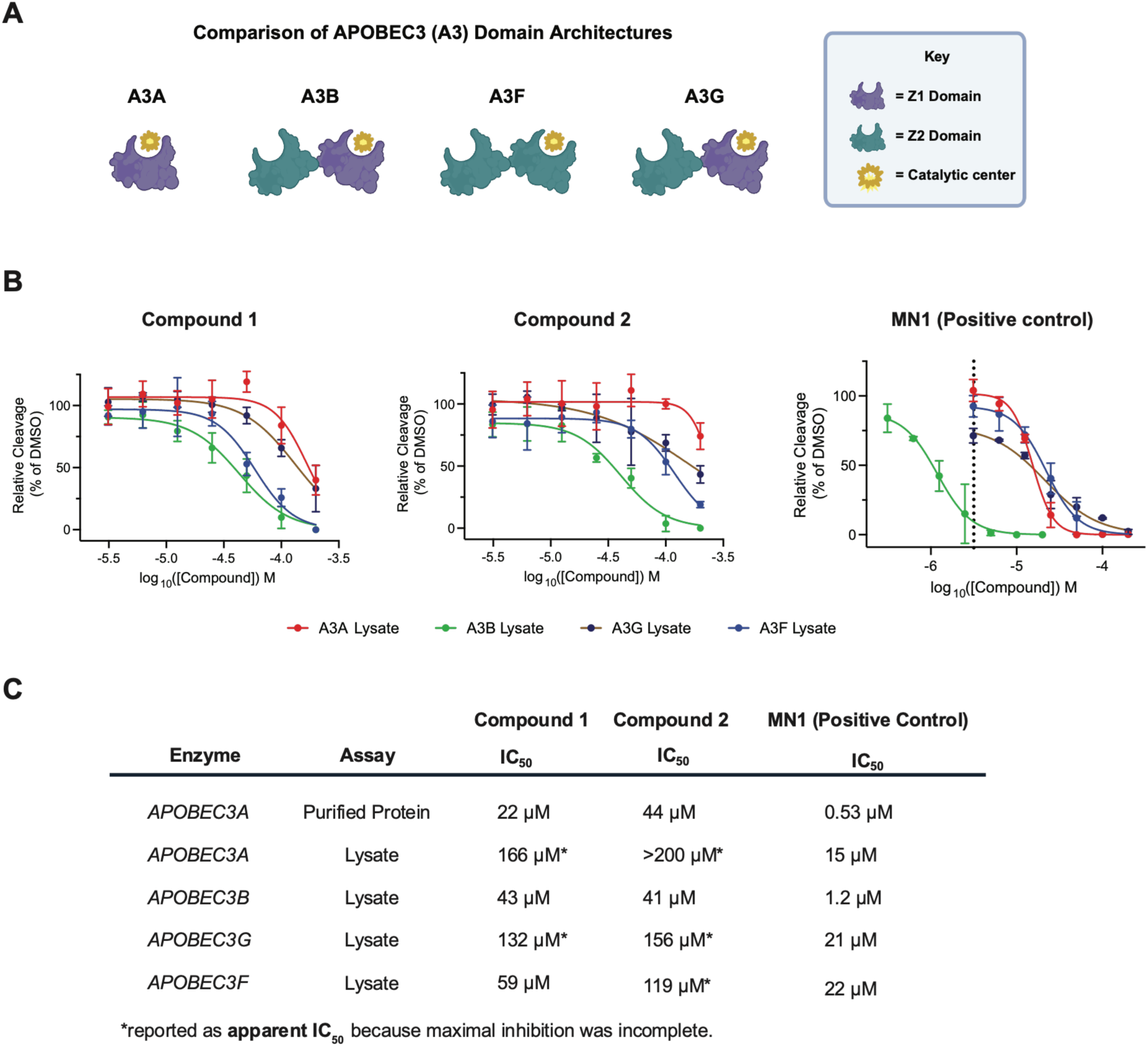
Compound dose-response analysis across APOBEC3 lysate-based deaminase assays (A) APOBEC3 (A3) domain architectures of the enzymes evaluated. (B) Dose-response curves for Compounds 1 and 2 and the positive control MN1 in A3A, A3B, A3F, and A3G lysate-based deaminase assays. Curves were fit using a four-parameter logistic model with the bottom constrained to 0% relative cleavage. (C) Summary of IC₅₀ values for Compounds 1 and 2 and MN1. IC₅₀ values are reported for purified A3A and lysate-based assays. Apparent IC₅₀ values (denoted by *) indicate that maximal inhibition was not achieved within the concentration range tested.

## Discussion

In this study, we identified APOBEC3 family inhibitors using a fluorescence polarization-based disulfide fragment screen targeting the A3A–ssDNA complex. Orthogonal LC–MS analyses revealed that only a subset of hits covalently modified C64, while subsequent studies demonstrated that cysteine tethering is not required for inhibition by the lead compounds, Compounds 1 and 2. Both compounds inhibited the deaminase activity of A3A, A3B, A3G, and A3F without detectable inhibition of Class I/II HDACs.

These findings establish an integrated workflow for APOBEC ligand discovery and identify the dichlorophenol scaffold as a promising starting point for APOBEC-directed chemical probes. The broad activity of this scaffold suggests engagement of conserved features shared across APOBEC3 enzymes, providing a foundation for future medicinal chemistry to improve potency and engineer family-member selectivity. Such compounds will serve as valuable tools for studying APOBEC biology and evaluating the therapeutic potential of APOBEC inhibition.

## METHODS

### APOBEC3A^E72A^ Protein Purification

A3A^E72A^ was expressed and purified as previously described with minor modifications.^35^ Detailed expression and purification procedures are provided in the Supporting Information. The same procedure was used to purify the A3A^E72A/C64S^ variant. Protein purity (>95%) was confirmed by SDS-PAGE (Figure S2).

### Fluorescence Polarization Protein Titration Assays

Fluorescence polarization (FP) binding assays were performed using purified A3A^E72A^ and a Cy5-labeled ssDNA substrate as previously described with minor modifications.^33^ Detailed assay conditions are provided in the Supporting Information. Apparent dissociation constants (K_d_) were determined by nonlinear regression using GraphPad Prism 10.0.

### Fluorescence Polarization Disulfide Fragment Screening

High-throughput fluorescence polarization (FP) screening was performed to identify disulfide fragments that disrupted A3A^E72A^ binding to a Cy5-labeled ssDNA tracer. Assays were performed using 3.2 μM A3A^E72A^ (EC₉₀ of tracer binding) and 5 nM Cy5-labeled ssDNA in optimized assay buffer. Disulfide fragments were screened at a final concentration of 200 μM in 384-well plates. After 3 h incubation at room temperature, fluorescence polarization was measured using an EnVision Xcite plate reader. Hits were defined as compounds producing polarization values ≥3 standard deviations below the no-compound control. Assay performance was assessed by the Z′ factor, which was 0.7. Detailed screening conditions are provided in the Supporting Information.

### Mass Spectrometry Tethering Assays

Mass spectrometry (MS) tethering assays were performed using purified A3A^E72A^ to evaluate covalent disulfide fragment binding. Disulfide fragments were screened at a final concentration of 200 μM and incubated with protein for 3 h at room temperature. Samples were analyzed by LC–MS, and percent tethering was determined from deconvoluted mass spectra as the fraction of compound-bound protein relative to total protein. Site specificity was assessed using the A3A^E72A/C64S^ variant. Detailed assay conditions are provided in the Supporting Information.

### A3A^WT^ Protein Purification

A3A^WT^ was expressed and purified from Expi293F cells as previously described with minor modifications.^49^ Detailed expression and purification procedures are provided in the Supporting Information. The same procedure was used to purify A3A^C64S^. Protein expression and purity were confirmed by SDS-PAGE and anti-c-Myc immunoblotting, and protein concentrations were determined by BCA assay.

### Gel-Based DNA Deaminase Assays with Purified Protein

Gel-based DNA deaminase assays were performed as previously described with minor modifications. ^37^ Purified A3A^WT^-Myc-His was preincubated with compounds prior to initiation of the reaction using a fluorescent ssDNA substrate. Cleavage products were resolved by denaturing PAGE and quantified by fluorescence imaging. Relative deaminase activity was calculated by normalizing substrate cleavage to DMSO-treated controls, and IC₅₀ values were determined by nonlinear regression using GraphPad Prism 10.0. The same assay conditions were used for A3A^C64S^. Detailed assay conditions are provided in the Supporting Information.

### Gel-Based DNA Deaminase Assays with Cell Lysates

HEK293T lysates expressing A3A^WT^-Myc-His, A3B^WT^-Myc-His, A3G^WT^-Myc-His, or A3F^WT^- Myc-His were prepared as previously described.³⁹ Gel-based DNA deaminase assays were performed using optimized lysate inputs and enzyme-specific ssDNA substrates as described above. Dose-response experiments were performed using seven-point two-fold serial dilutions of compounds beginning at 200 μM. Cleavage products were analyzed by denaturing PAGE, and deaminase activity was quantified as described for purified protein. A3B^C247S^ lysates were prepared and assayed under identical conditions. Detailed procedures are provided in the Supporting Information.

### Molecular Docking

Molecular docking simulations were carried out using Molecular Operating Environment (MOE 2024.0601).^50^ The three-dimensional crystal structure of human APOBEC3A^E72A^ (5SWW) was downloaded from Protein Data Bank (PDB). Alanine-72 was mutated back to glutamate within Pymol version 2.4.1 to generate the wild type protein. The resulting protein structure was prepared for docking within MOE using default parameters.

Compounds 1 and 2 were separately docked within MOE, using Triangle Matcher scored by London dG with 30 poses and refined by induced fit to identify the top 5 poses. Ligand interactions were determined within MOE for the top pose for each compound. Images of the docked structures were produced in Pymol.

## ASSOCIATED CONTENT

### Supporting Information

The following files are available free of charge.

Supplementary methods, supplementary results, supplementary figures, chemical synthesis procedures, compound characterization, ^1^H and ^13^C NMR spectra, and supplementary references (PDF).

### Author Contributions

J.S.B. and H.R.A. contributed equally to this work. J.S.B. and H.R.A. designed and performed the biochemical experiments and analyzed the data. W.L. assisted with bacterial and mammalian protein expression. J.L.R. and M.R.A. contributed to the design of the fluorescence polarization assay and compound screening. R.B. contributed to the coordination and analysis of the chemical synthesis, and C.M.S. contributed to the analysis of the chemical characterization data. J.M.L.O. conceptualized and supervised the project. The manuscript was written through contributions of all authors. All authors have given approval to the final version of the manuscript.

### Funding Sources

This work was supported by the National Cancer Institute of the National Institutes of Health under Award Number U54CA224081.

### Notes

The authors declare no competing financial interests.

## ACKNOWLEDGMENT

We thank Pharmaron for the synthesis, characterization, and acquisition of NMR spectra for the compounds used in this study. We also thank the Small Molecule Discovery Center (SMDC) at the University of California, San Francisco for assistance with compound screening.

## ABBREVIATIONS

A3A: APOBEC3A
A3B: APOBEC3B
FP: fluorescence polarization
ssDNA: single-stranded DNA
UDG: uracil-DNA glycosylase
LC–MS: liquid chromatography–mass spectrometry
DCP: dichlorophenol.

