## Supplementary Information for "Fluorescence polarization-based fragment screen identifies inhibitors of APOBEC3A and APOBEC3B"

‡ Equal Contribution

**Table of Contents**

Supplementary Methods……………………………………………………………………S2

Supplementary Results…………………………………………………………………..S9

Chemical Synthesis……………………………………………………………………..S23

Compound Characterization………………………………………………………………S33

Supplementary References………………………………………………………………..S45

### Supplementary Methods

#### APOBEC3A$^{E72A}$ Protein Purification

A3A$^{E72A}$ was expressed and purified as previously described, with the following modifications. Human APOBEC3A containing the catalytic substitution E72A was expressed in *Escherichia coli* BL21(DE3) Star cells from a pET-28a(+) vector encoding an N-terminal $His_6$-GST fusion protein. Cultures were supplemented with 100 µM $ZnCl_2$ and induced with 0.5 mM IPTG, followed by overnight expression at 18 °C. Cell pellets were resuspended in lysis buffer containing 20 mM Tris-HCl (pH 8.0), 500 mM NaCl, and 10 mM β-mercaptoethanol supplemented with EDTA-free protease inhibitor cocktail (Roche) and lysozyme. Following sonication, clarified lysates were applied to a HisTrap HP column (Cytiva), and bound protein was eluted using a linear gradient to 400 mM imidazole. Fractions containing A3A were pooled and dialyzed overnight at 4 °C against 20 mM Tris-HCl (pH 8.0), 500 mM NaCl, and 10 mM β-mercaptoethanol. The $His_6$-GST tag was removed by incubation with HRV 3C protease (Fisher Scientific) at a 1:200 (w/w) protease-to-protein ratio for 1 h at 4 °C. The cleavage reaction was applied to a GSTrap 4B column (Cytiva) equilibrated in the dialysis buffer, allowing GST-tagged species and uncleaved fusion protein to be retained while cleaved A3A was collected in the flow-through. Bound proteins were subsequently eluted using a linear gradient of reduced glutathione to a final concentration of 10 mM. Final purification was achieved by size-exclusion chromatography on a Superdex 75 column (Cytiva) equilibrated in 20 mM Tris-HCl (pH 7.4), 200 mM NaCl, and 0.5 mM TCEP. To improve protein expression and solubility, residual N-terminal glycine-asparagine residues were removed from the construct. The same procedure was used for purification of the A3A$^{E72A/C64S}$ variant. Protein purity (>95%) was confirmed by SDS-PAGE and Coomassie staining. (Figure S2).

**Fluorescence Polarization Protein Titration Assays**

Fluorescence polarization (FP) protein dose–response assays were performed using purified A3A$^{E72A}$. Assays were conducted in buffer containing 50 mM HEPES (pH 7.5), 100 mM NaCl, 2.0 mM tris(2-carboxyethyl)phosphine hydrochloride (TCEP-HCl), and 4 mM CHAPS, with conditions adapted from previously reported fluorescence polarization assays.[1] For pH-dependent studies (pH 5.0–6.5), 50 mM MES was substituted for HEPES. A Cy5-labeled single-stranded DNA tracer (5′-AAAAAAATCGGGAAA-3′; Integrated DNA Technologies; MW = 5183.8 g/mol) was used at a final concentration of 5 nM. Protein dose–response experiments were performed using a 24-point 1:1 serial dilution of A3A$^{E72A}$, beginning at 100 μM and yielding a highest final assay concentration of 50 μM. Assays were assembled in black low-volume 384-well plates (Corning 4514) in a final volume of 10 μL and incubated at room temperature for 30 min prior to measurement. Fluorescence polarization was measured using an EnVision Xcite multimode plate reader (PerkinElmer; Model 2105). Data were analyzed in GraphPad Prism 10.0 using a four-parameter nonlinear regression model (variable slope) to determine apparent dissociation constants ($K_d$).

**Fluorescence Polarization Disulfide Fragment Screening**

High-throughput fluorescence polarization (FP) screening was performed to identify disulfide fragments that disrupt A3A$^{E72A}$ binding to a Cy5-labeled ssDNA tracer. Assays were conducted under conditions determined from fluorescence polarization protein titration assays, with A3A$^{E72A}$ used at a final concentration of 3.2 μM, corresponding to the $EC_{90}$ of tracer binding. Screening reactions were performed in assay buffer containing 50 mM HEPES (pH 7.5), 100 mM NaCl, 0.5 mM tris(2-carboxyethyl)phosphine hydrochloride (TCEP-HCl), and 4 mM CHAPS. The Cy5-labeled tracer was used at a final concentration of 5 nM. Compounds were transferred from 50 mM stock solutions using an Echo acoustic liquid handler (Labcyte), dispensing 40 nL per well to achieve a final compound concentration of 200 μM. Reactions were assembled in a final volume of 10 μL using a Dragonfly Discovery liquid handler (SPT Labtech). Control wells contained either all assay components except protein (negative control) or complete reaction mixtures lacking compound (positive control). Plates were incubated at room temperature for 3 h prior to measurement. Fluorescence polarization was measured using an EnVision Xcite multimode plate reader (PerkinElmer; Model 2105). Hits were defined as compounds producing polarization values ≥3 standard deviations below the mean of the no-compound control wells. Assay performance was evaluated using the Z′ factor, with a value of 0.7 indicating robust separation between positive and negative controls.

**Mass Spectrometry Tethering Assays**

Mass spectrometry (MS) tethering assays were performed using purified A3A$^{E72A}$ at a final concentration of 1 μM in buffer containing 10 mM Tris-HCl (pH 8.3) and 0.5 mM tris(2-carboxyethyl)phosphine (TCEP). Disulfide fragments were transferred from library plates using an Echo acoustic liquid handler (Labcyte), dispensing 100 nL of 50 mM stock solutions to achieve a final compound concentration of 200 μM. Reactions were initiated by addition of protein master mix to a final volume of 25 μL per well and incubated at room temperature for 3 h. For iodoacetamide labeling experiments, purified A3A$^{E72A}$ (1 μM) was incubated with either 0 or 1 mM iodoacetamide under identical buffer conditions for 3 h at room temperature. Samples were analyzed by liquid chromatography–mass spectrometry using a Waters Xevo Q-tof mass spectrometer. Percent tethering was determined from deconvoluted mass spectra as the fraction of compound-bound protein relative to total protein (bound plus unbound). Site specificity was assessed using the A3A$^{E72A/C64S}$ variant under identical assay conditions.

**A3A$^{WT}$ Protein Purification**

A3A$^{WT}$ was expressed and purified based on previously reported methods, with the following modifications.[2] Expi293F cells were transiently transfected with pcDNA3.1-A3A$^{WT}$-Myc-His using the ExpiFectamine 293 Expression System (Thermo Fisher Scientific) according to the manufacturer's instructions. Cells were harvested 5 d post-transfection and lysed in buffer containing 25 mM HEPES (pH 7.5), 150 mM NaCl, 0.5% Triton X-100, 1 mM $MgCl_2$, 10% glycerol, EDTA-free protease inhibitor cocktail (cOmplete Mini, EDTA-free; Roche), and 50 μg/mL RNase A (Qiagen). Clarified lysates were adjusted to 0.8 M NaCl and incubated with Ni-NTA agarose resin (Qiagen) overnight at 4 °C. The resin was washed with 50 mM Tris-HCl (pH 8.0), 300 mM NaCl, 10% glycerol, 0.5% Triton X-100, and 50 mM imidazole, and bound protein was eluted with the same buffer containing 150 mM imidazole. Eluted fractions were analyzed by SDS-PAGE and Coomassie staining, and protein expression was confirmed by immunoblotting using an anti-c-Myc antibody (9E11; antibodies.com). Protein concentrations were determined using a BCA assay. Where indicated, samples were dialyzed into storage buffer consisting of 50 mM Tris-HCl (pH 8.0), 300 mM NaCl, 10% glycerol, and 0.5% Triton X-100 prior to downstream assays. The same procedure was used for purification of A3A$^{C64S}$.

**Gel-Based DNA Deaminase Assays with Purified Protein**

Gel-based DNA deaminase assays were performed as previously described, with the following modifications.[3] Purified A3A$^{WT}$-Myc-His was diluted to a final concentration of 100 nM in assay buffer containing 50 mM Tris-HCl (pH 7.5), 30 mM NaCl, and 0.5 mM TCEP. Protein was preincubated with compounds for 1 h at room temperature in a reaction volume of 45 μL. For dose–response experiments, compounds were tested in a seven-point two-fold serial dilution series beginning at 200 μM. Reactions were initiated by addition of a 5 μL master mix containing single-stranded DNA substrate and uracil DNA glycosylase (UDG), yielding final concentrations of 0.4 μM ssDNA and 1.5 U UDG in a total reaction volume of 50 μL. The DNA substrate sequence was 5′-(6-FAM)-GCAAGCTGGTCGGAAAAATGA-3′ (Integrated DNA Technologies; MW = 7065.8 g/mol). An uracil-containing control substrate was included as a positive control for UDG-mediated cleavage. Reactions were incubated at 37 °C for 1 h and terminated by addition of NaOH to a final concentration of 0.25 N, followed by incubation at 95 °C for 40 min. Samples were mixed with 2× Novex TBE-Urea Sample Buffer (Thermo Fisher Scientific), heated at 95 °C, and resolved on 15% denaturing TBE-urea polyacrylamide gels (Novex). Gels were imaged using a ChemiDoc MP imaging system (Bio-Rad). Band intensities were quantified using ImageJ. Fractional cleavage was calculated as the intensity of the cleaved product divided by the sum of cleaved and uncleaved substrate intensities. For inhibitor studies, cleavage values were normalized to DMSO-treated controls and reported as relative deaminase activity. Data was analyzed in GraphPad Prism 10.0 using a four-parameter nonlinear regression model (variable slope) to determine $IC_{50}$ values. The same assay conditions were used for A3A$^{C64S}$.

Counter screening against uracil DNA glycosylase (UDG) was performed using a uracil-containing substrate, 5′-(6-FAM)-GCAAGCTGGTUGGAAAAATGA-3′ (Integrated DNA

Technologies; MW = 7066.8 g/mol). Reactions were assembled as described above for the gel-based DNA deaminase assay, except that compounds were tested at a final concentration of 100 µM and the uracil-containing substrate was substituted for the cytosine-containing substrate. UDG-mediated cleavage was quantified as described above.

**Gel-Based DNA Deaminase Assays with Cell Lysates**

HEK293T lysates expressing A3A$^{WT}$-Myc-His, A3B$^{WT}$-Myc-His , A3G$^{WT}$-Myc-His, or A3F$^{WT}$-Myc-His were prepared as previously described.[3] Cells were transiently transfected with the indicated expression constructs and harvested 24 h post-transfection for A3A or 48 h post-transfection for A3B, A3G, and A3F. Cells were lysed in buffer containing 25 mM HEPES (pH 7.9), 150 mM NaCl, 1 mM $MgCl_2$, 10% glycerol, and 0.5% Triton X-100. Clarified lysates were stored at −80 °C until use.

Gel-based DNA deaminase assays were performed as described above for purified protein, with the following modifications. Reactions were conducted in buffer containing 50 mM Tris-HCl (pH 7.5), 30 mM NaCl, 0.5 mM TCEP, 0.25 mM $ZnCl_2$, and 10 mM EDTA. Lysate input was optimized to achieve partial substrate turnover within the linear range of the assay. For dose–response experiments, compounds were tested in seven-point two-fold serial dilution series beginning at 200 µM. No compound preincubation step was performed. Enzyme-specific ssDNA substrates were selected based on the preferred deamination sequence of each APOBEC enzyme as previously described.[4,5] The A3G substrate was 5′-(6-FAM)-ATTATTATTATTATTCCCGATTATTTATTTATTTATTTATTT-3′ (Integrated DNA Technologies; MW = 12,873.3 g/mol), and the A3F substrate was 5′-(6-FAM)-ATTTATATTATTTATTCATATTTATATTTA-3′ (Integrated DNA Technologies; MW = 9,218.9 g/mol).Reactions were incubated at 37 °C for 4 h prior to NaOH cleavage, gel electrophoresis, and quantification as described above. $A3B^{C247S}$ lysates were prepared and assayed under the same conditions as $A3B^{WT}$. Data were analyzed as described above.

**HDAC Counter-Screen Assay**

HDAC inhibition was evaluated using the HDAC-Glo I/II Assay (Promega) according to the manufacturer's instructions with modifications for HEK nuclear lysates. Nuclear lysates were prepared from HEK cells grown to approximately 90% confluency by detergent-based isolation of nuclei followed by extraction in lysis buffer (50 mM Tris-HCl, 150 mM NaCl, 10% glycerol, 0.5% Triton X-100) containing Halt Protease Inhibitor Cocktail (Thermo Scientific). Protein concentration was determined by bicinchoninic acid (BCA) assay. To determine the appropriate enzyme concentration, nuclear lysates were evaluated in a six-point two-fold serial dilution beginning at 100 μg/mL. A final lysate concentration of 12.5 μg/mL during the compound incubation was selected because it fell within the linear range of the assay. For inhibitor studies, Compound 1, Compound 2, and MN1 were tested in 10-point two-fold serial dilution series beginning at 200 μM (final incubation concentration). Trichostatin A (TSA) was included as a positive control and evaluated in a 10-point two-fold serial dilution beginning at 50 nM (final incubation concentration). Compound solutions (25 μL) were mixed with an equal volume of HEK nuclear lysate (25 μg/mL, 2× working concentration) and incubated for 30 min at room temperature. HDAC-Glo reagent (50 μL) was then added, and luminescence was measured after incubation according to the manufacturer's protocol. Data were analyzed in GraphPad Prism 10.0 using a four-parameter nonlinear regression model (variable slope).

### Supplementary Results

#### Optimization of Fluorescence Polarization Assay Conditions for A3A–ssDNA Binding

To establish conditions suitable for fluorescence polarization (FP)-based screening, a panel of fluorescently labeled ssDNA substrates containing the canonical 5′-TC-3′ motif was evaluated (Figure S1A). Variations in the nucleotides flanking the target cytosine were introduced to probe sequence-context effects on binding, as A3A has previously been reported to preferentially target YTCA motifs (Y = pyrimidine).[6,7] Despite these sequence variations, only modest differences in binding affinity were observed, indicating that recognition was primarily driven by the central TC motif. Based on its high affinity and prior structural validation, the sequence 5′-Cy5-AAAAAAATCGGGAAA-3′ was selected for subsequent studies.[7,8] Cy5 labeling was further favored because its red-shifted emission minimized fluorescence interference from disulfide fragments during screening.

The influence of pH on A3A–ssDNA binding was next evaluated across a range of pH 5.5–8.5 (Figure S1B). Binding affinity increased under acidic conditions and progressively decreased at higher pH values. Although maximal binding was observed at lower pH, pH 7.5 was selected for subsequent experiments because it provided a robust assay window while maintaining physiological relevance.

To further assess substrate specificity, binding was measured using substrates containing TC, TU, or TA target motifs (Figure S1C). A3A exhibited strong binding to TC-containing substrates, reduced binding to TU-containing substrates, and little to no detectable interaction with TA-containing substrates. These observations are consistent with previous reports demonstrating preferential recognition of cytosine-containing substrates and reduced affinity for uracil-containing reaction products.[9]

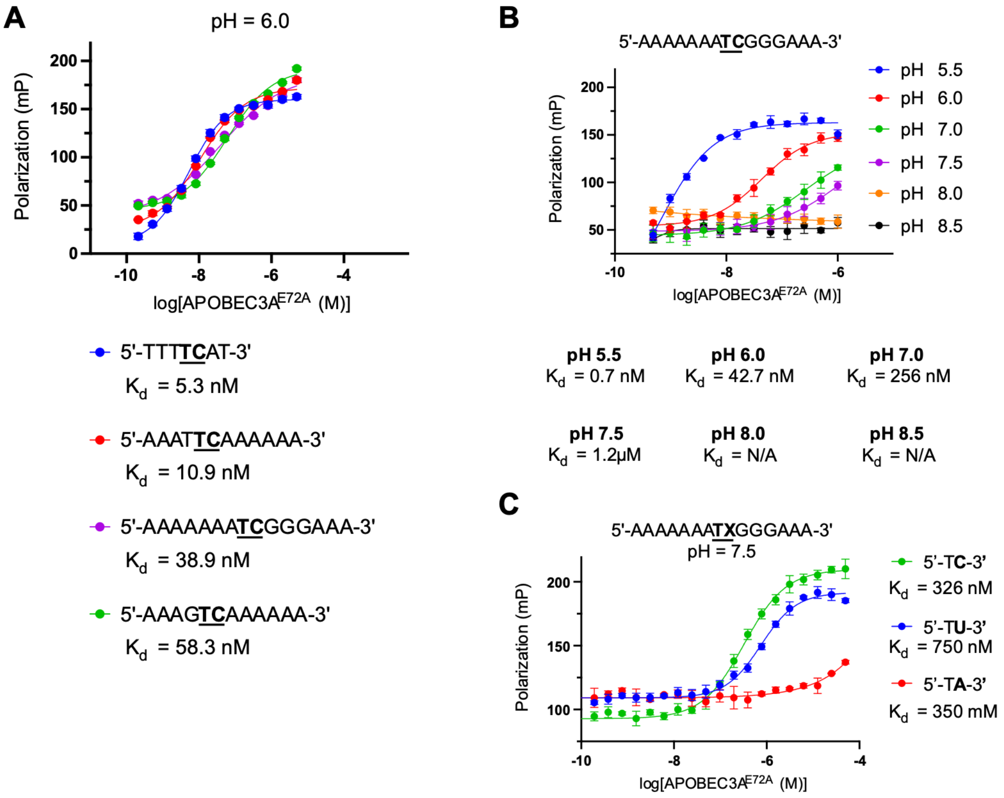


**Figure S.1** Optimization of fluorescence polarization assay conditions for A3A–ssDNA binding. (A) Binding of A3A$^{E72A}$ to ssDNA substrates containing the canonical 5′-TC-3′ motif embedded within distinct flanking sequence contexts. Variation in the surrounding nucleotides produced only modest effects on binding affinity. (B) Effect of pH on A3A$^{E72A}$–ssDNA binding. Binding affinity increased under acidic conditions and decreased at higher pH values. pH 7.5 was selected for subsequent screening experiments to balance assay performance with physiological relevance. (C) Influence of the target dinucleotide on A3A$^{E72A}$ binding. A3A$^{E72A}$ bound preferentially to TC-containing substrates, exhibited reduced affinity for TU-containing substrates, and showed minimal interaction with TA-containing substrates.

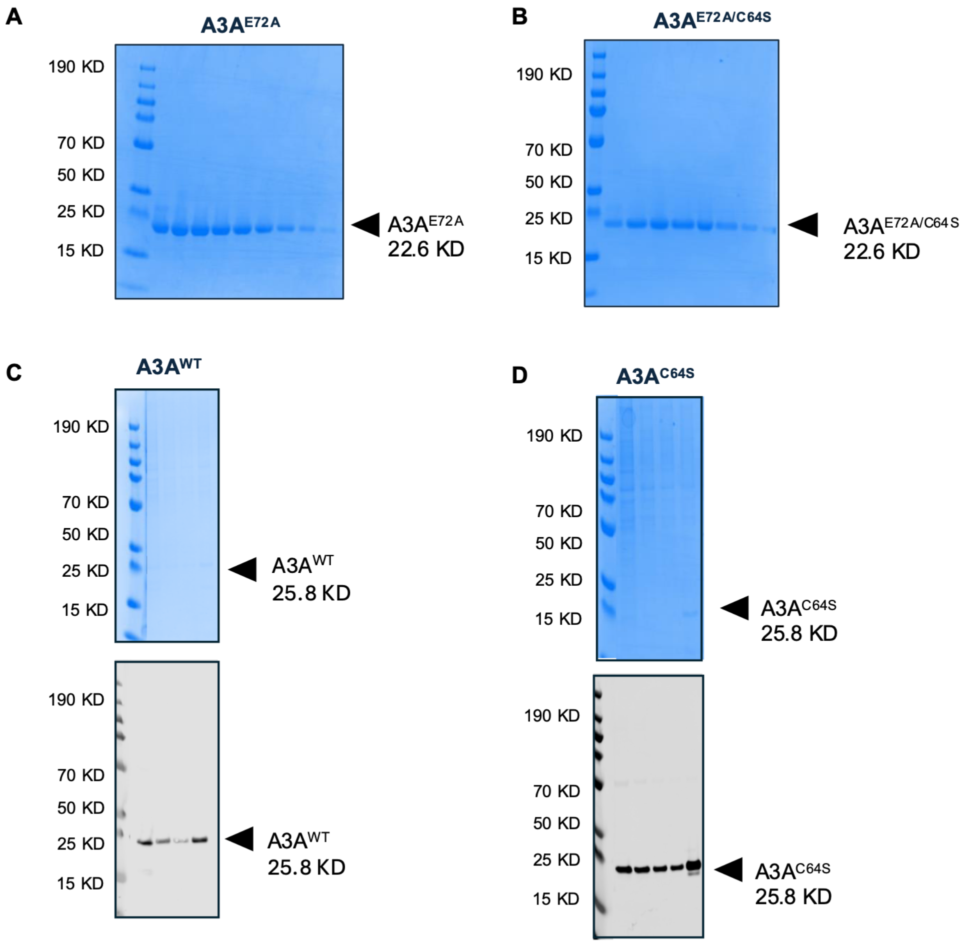


**Figure S.2** Purification of recombinant A3A variants used in biochemical assays. (A) Coomassie-stained SDS-PAGE gel of purified A3A$^{E72A}$ following affinity and size-exclusion chromatography. (B) Coomassie-stained SDS-PAGE gel of purified A3A$^{E72A/C64S}$ following affinity and size-exclusion chromatography. (C) Coomassie-stained SDS-PAGE gel and corresponding anti-Myc Western blot of purified A3A-WT following affinity chromatography. (D) Coomassie-stained SDS-PAGE gel and corresponding anti-Myc Western blot of purified A3A$^{C64S}$ following affinity chromatography. Arrowheads indicate the expected migration position of the indicated A3A proteins.

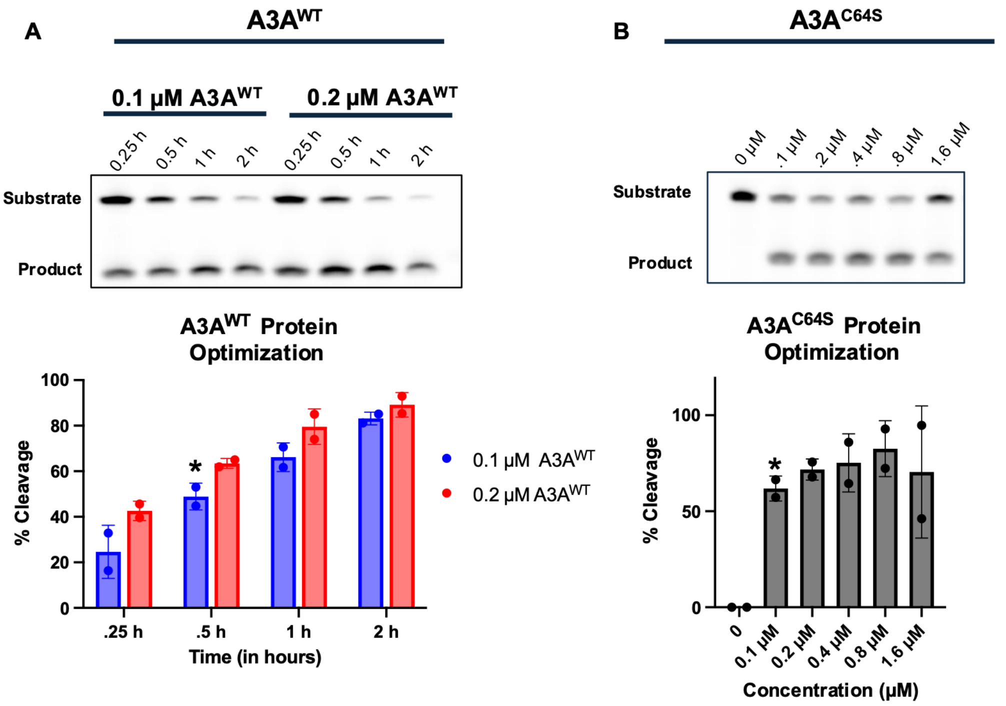


**Figure S.3** Optimization of A3A deaminase assay conditions. (A) Optimization of assay conditions for A3A$^{WT}$. Deaminase activity was monitored as a function of reaction time using 0.1 µM and 0.2 µM purified A3A$^{WT}$. Representative denaturing gels and corresponding quantification of substrate cleavage are shown. A reaction time of 30 min using 0.1 µM A3A$^{WT}$ produced partial substrate turnover within the dynamic range of the assay and was selected for subsequent inhibition studies. (B) Optimization of assay conditions for A3A$^{C64S}$. Using the 30 min reaction time established for A3A$^{WT}$, deaminase activity was evaluated across a range of A3A$^{C64S}$ concentrations. Representative denaturing gels and corresponding quantification of substrate cleavage are shown. A concentration of 0.1 µM A3A$^{C64S}$ produced substrate turnover comparable to that observed for A3A$^{WT}$ and was selected for subsequent inhibition studies. Asterisks indicate assay conditions used in subsequent experiments.

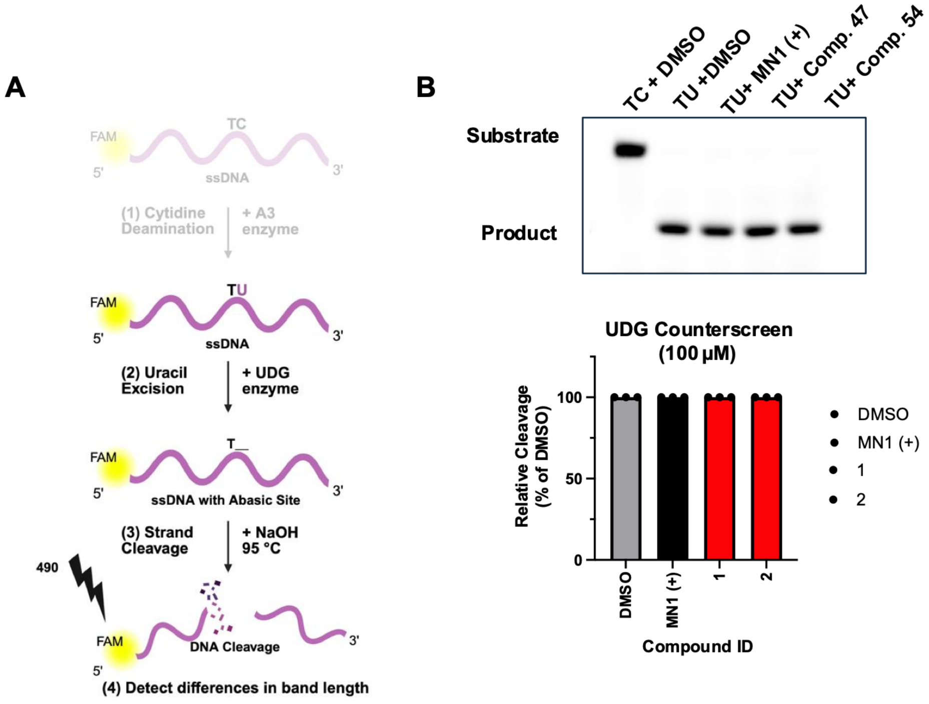


**Figure S.4** UDG counter-screen confirms A3A-specific inhibition. (A) Schematic of the UDG screen used to evaluate compound effects on the downstream coupling enzyme in the deaminase assay. In the absence of A3A, an uracil-containing ssDNA substrate is directly processed by UDG, enabling assessment of UDG activity independent of cytosine deamination. (B) Representative denaturing gel and corresponding quantification of UDG activity in the presence of the indicated compounds. Neither compound 1 nor compound 2 reduced UDG-mediated substrate cleavage under assay conditions. MN1 similarly showed no effect on UDG activity, indicating that inhibition observed in the deaminase assay arises from inhibition of A3A rather than interference with the downstream UDG-coupled readout.

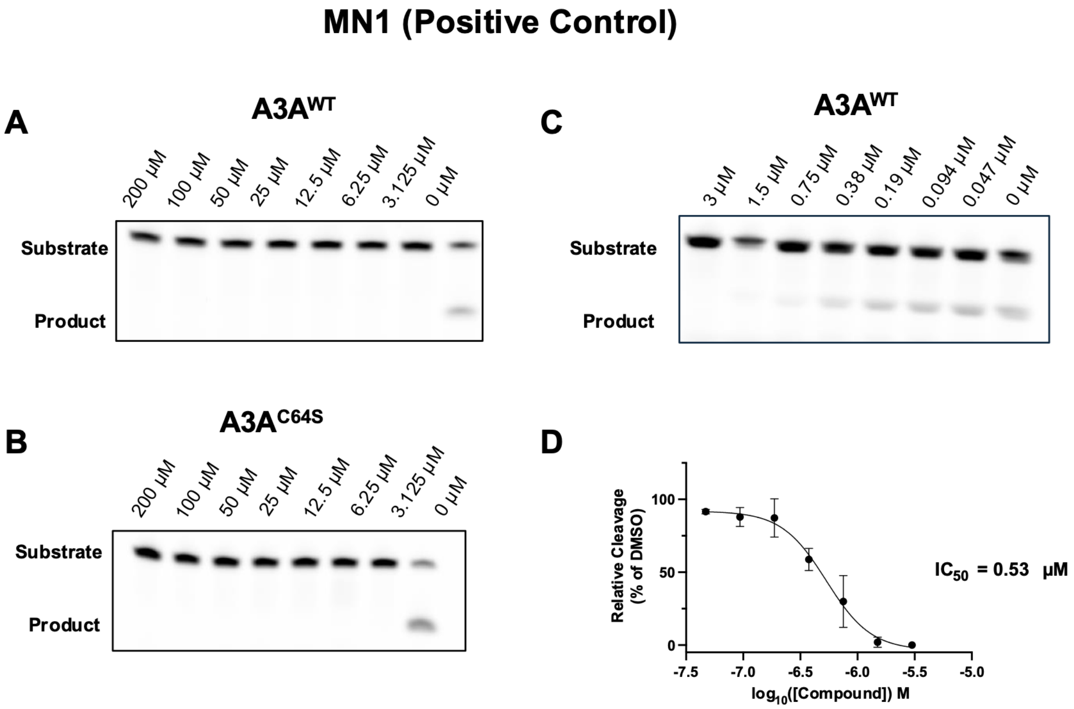


**Figure S.5** MN1 positive-control inhibition of A3A$^{WT}$ and A3A$^{C64S}$. (A-B) MN1 dose-response analysis in A3A$^{WT}$ and A3A$^{C64S}$ under the assay conditions used for compound testing. MN1 inhibited both proteins, demonstrating that the assay can detect inhibitory effects in either background and would reveal differences between WT and C64S if present. (C-D) Expanded dose-response analysis of MN1 against A3A$^{WT}$, yielding an apparent $IC_{50}$ of 0.53 μM.

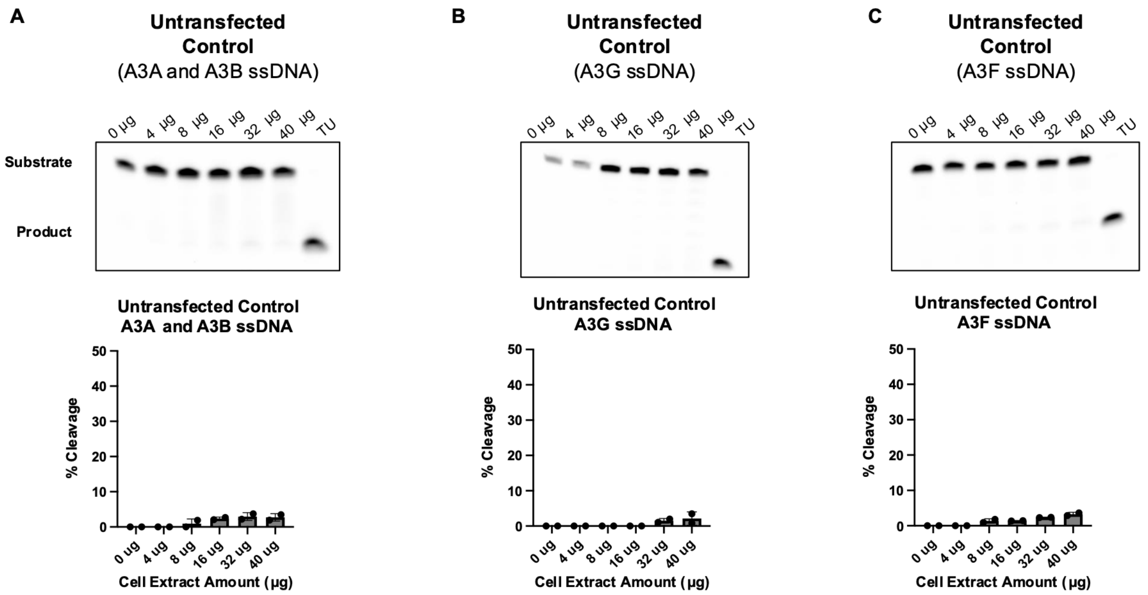


**Figure S.6** Untransfected Cell Lysate Control Demonstrates Negligible ssDNA Cleavage. Representative denaturing gels and quantification of ssDNA cleavage following incubation with increasing amounts of untransfected cell lysate (0–40 µg). Following the 0–40 µg lysate titration, a TU sequence was included as a positive control to indicate the expected migration position of the cleaved product band. (A) A3A/A3B ssDNA substrate. (B) A3G ssDNA substrate. (C) A3F ssDNA substrate. Across all three substrates, untransfected cell lysate produced negligible cleavage, with <4% cleavage observed even at the highest lysate concentration (40 µg), confirming that endogenous cellular nucleases do not contribute appreciably to the ssDNA cleavage observed in transfected lysates**.**

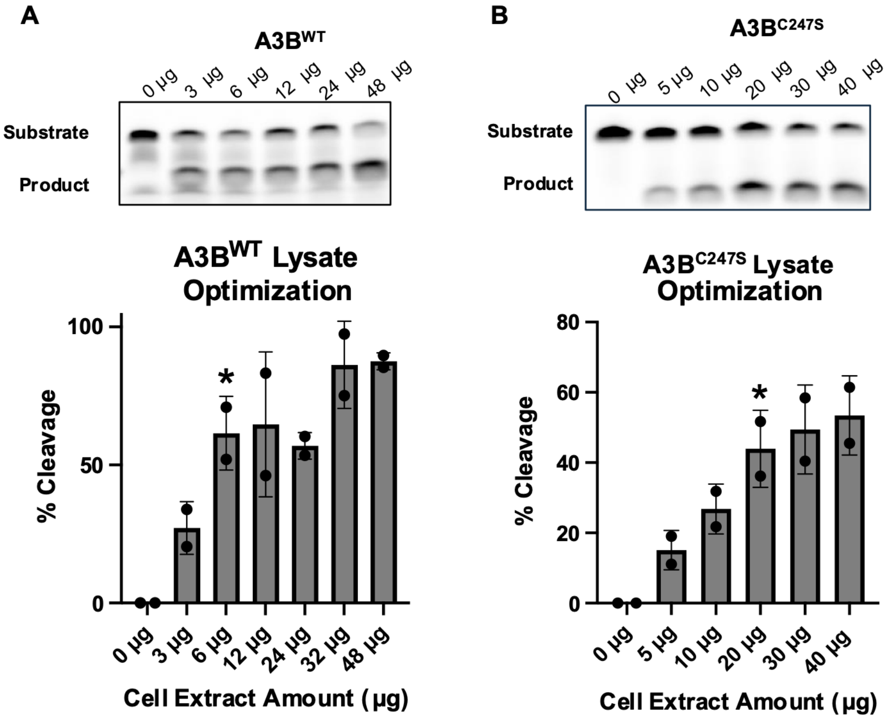


**Figure S.7** Optimization of A3B Lysate Deaminase Assay Conditions (A) Optimization of assay conditions for $A3B^{WT}$ lysate. Increasing amounts of lysate were evaluated to identify conditions that produced partial substrate turnover within the dynamic range of the assay. Representative denaturing gels and quantification of substrate cleavage are shown. A lysate input of 6 µg was selected for subsequent inhibition studies (*). (B) Optimization of assay conditions for $A3B^{C247S}$ lysate. Lysate input was varied to achieve substrate turnover comparable to that observed for $A3B^{WT}$. Representative denaturing gels and corresponding quantification are shown. A lysate input of 20 µg was selected for subsequent inhibition studies (*).

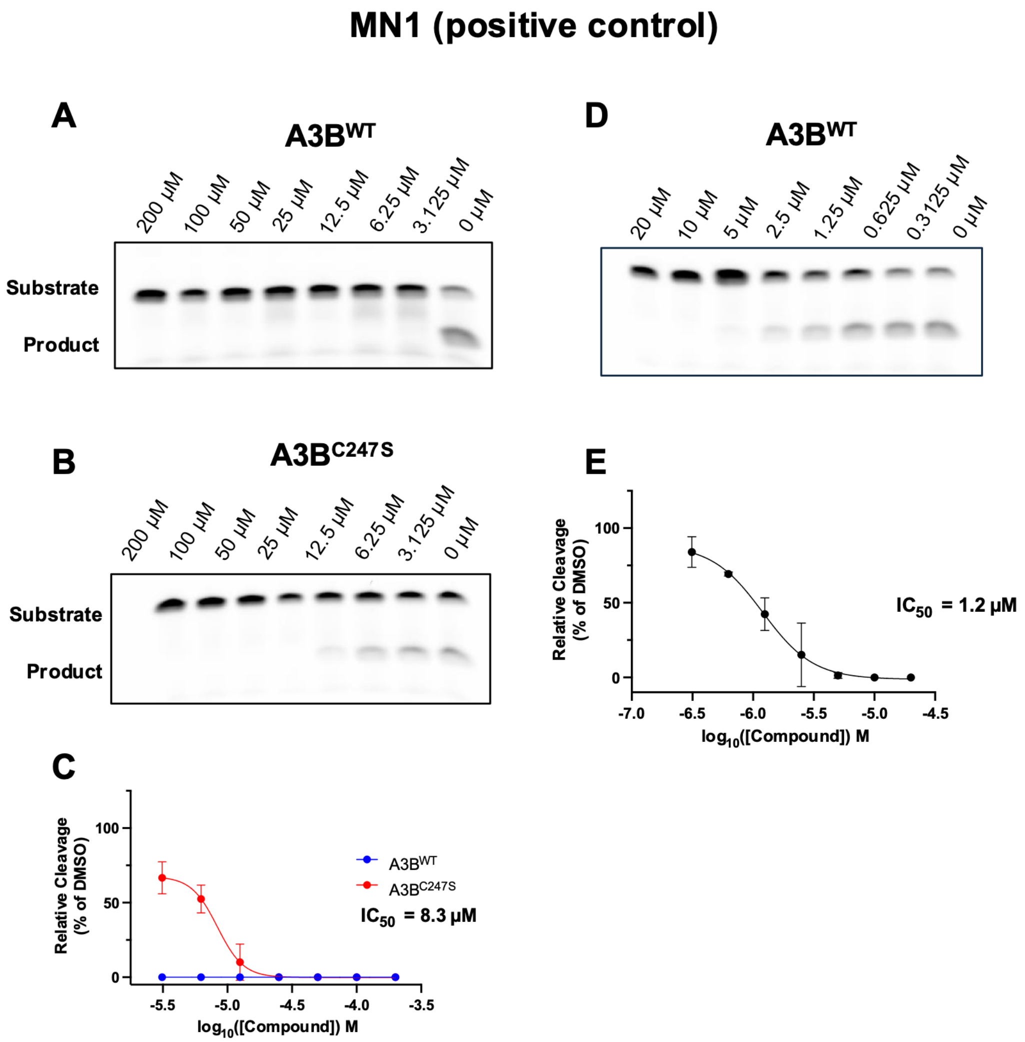


**Figure S.8** Positive-control inhibition establishes assay sensitivity in A3B$^{WT}$ and A3B$^{C247S}$ lysates (A) Representative dose-response inhibition of A3B$^{WT}$ lysate by the positive-control inhibitor MN1. (B) Representative dose-response inhibition of A3B$^{C247S}$ lysate by MN1. **(C)** Quantification of MN1-mediated inhibition in A3B$^{WT}$ and A3B$^{C247S}$ lysates. Although modest differences in apparent potency were observed between protein preparations, MN1 produced measurable inhibition in both backgrounds, indicating that the observed shifts likely reflect assay-dependent variation. An apparent $IC_{50}$ of 8.3 μM was determined for A3B$^{C247S}$ under these assay conditions. (D) Expanded dose-response analysis of MN1 in A3B$^{WT}$ lysate showing a representative gel. (E) Quantification of the experiment shown in (D). An apparent $IC_{50}$ of 1.2 μM was determined for A3B$^{WT}$ under these assay conditions.

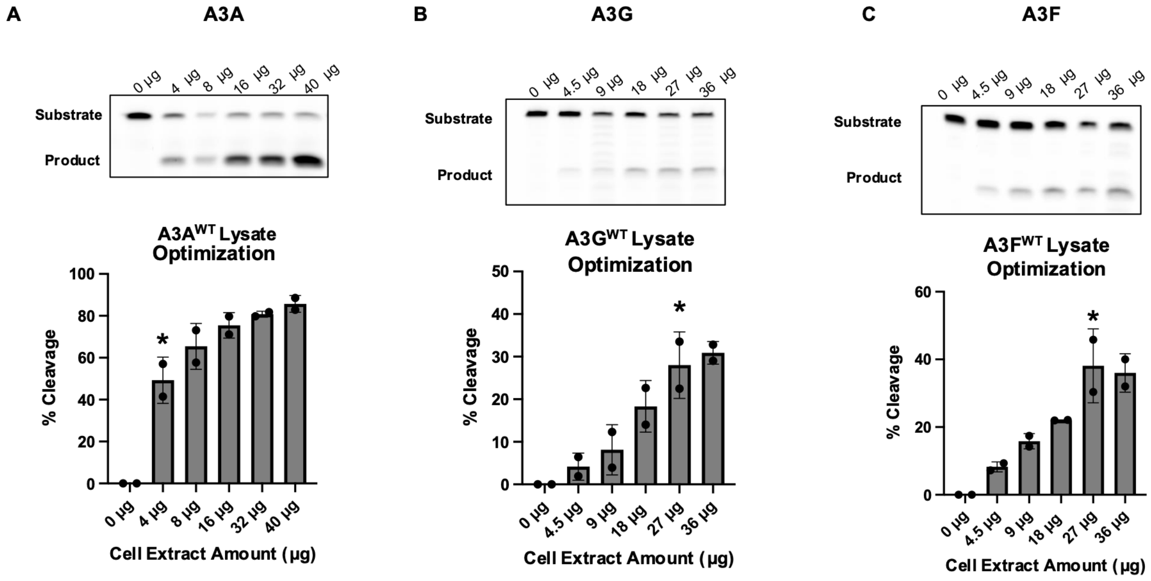


**Figure S.9** Optimization of Lysate Deaminase Assay Conditions for A3A, A3G, and A3F Increasing amounts of lysate were evaluated to identify conditions that produced partial substrate turnover within the dynamic range of the assay. Representative denaturing gels and corresponding quantification of substrate cleavage are shown. (A) A3A lysate optimization. A lysate input of 4 μg was selected for subsequent inhibition studies (*). (B) A3G lysate optimization. A lysate input of 27 μg was selected for subsequent inhibition studies (*). (C) A3F lysate optimization. A lysate input of 27 μg was selected for subsequent inhibition studies (*).

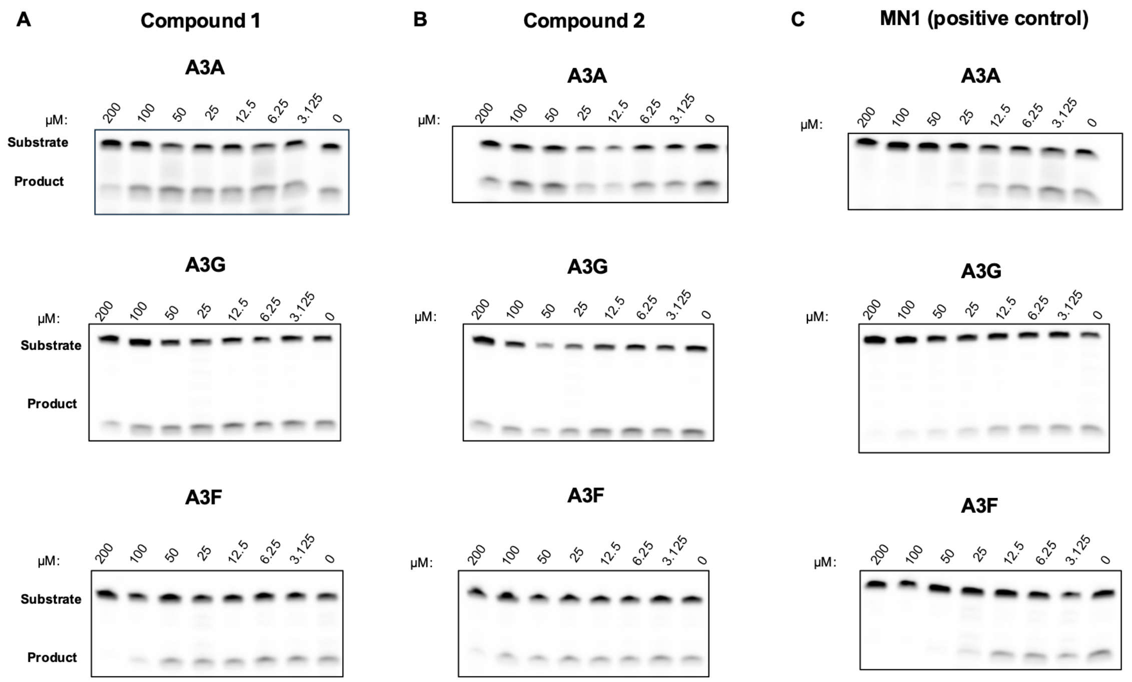


**Figure S.10** Representative denaturing gels from compound dose-response lysate assays. Representative gels corresponding to the quantified dose-response curves in Figure 7 are shown for A3A, A3G, and A3F lysate deaminase assays. Compounds tested were (A) Compound 1, (B) Compound 2, and (C) MN1 (positive control).

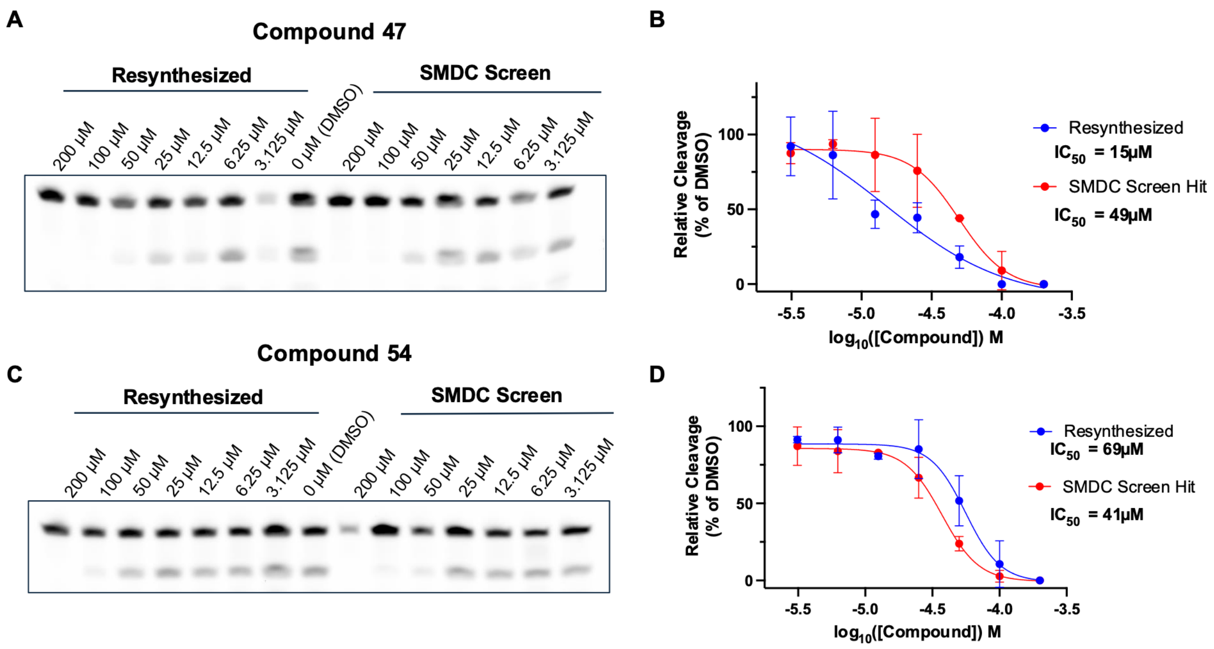


**Figure S.11** Validation of Small Molecule Discovery Center (SMDC) screening hits following compound resynthesis. Independently resynthesized compounds were retested in the A3A-C64S deaminase assay to confirm the reproducibility of the inhibitory activity observed during the primary screen. (A) Representative deaminase assay comparing independently resynthesized compound 47 with the corresponding compound from the SMDC screening library across the indicated concentrations. (B) Quantification of the assay shown in (A), demonstrating comparable dose-dependent inhibition and $IC_{50}$ values between the resynthesized compound and the original screening hit. (C) Representative deaminase assay comparing independently resynthesized compound 54 with the corresponding SMDC screening library compound. (D) Quantification of the assay shown in (C), demonstrating comparable dose-dependent inhibition and $IC_{50}$ values between the resynthesized compound and the original screening hit. These results confirm that the inhibitory activity of compounds 47 and 54 is reproducible following independent chemical synthesis.

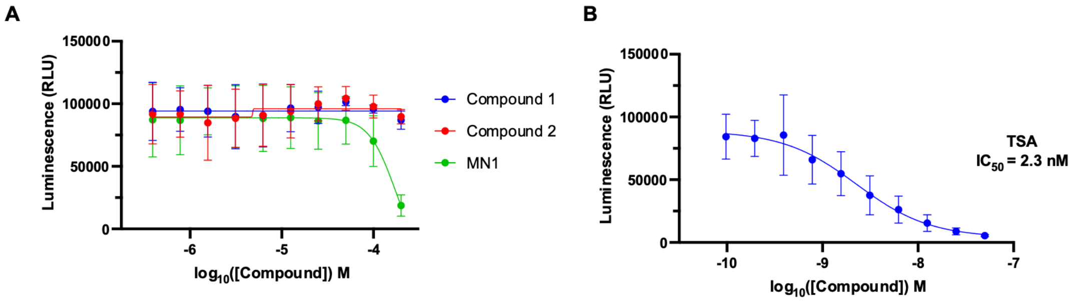


**Figure S.12** HDAC counter-screen demonstrates that Compound 1 and Compound 2 do not inhibit Class I/II HDAC activity. (A) HDAC-Glo I/II assay using HEK nuclear lysates incubated with Compound 1, Compound 2, or MN1 over a 10-point two-fold concentration series with a highest concentration of 200 µM. Compound 1 and Compound 2 did not reduce HDAC activity across the concentration range tested. MN1 exhibited inhibition only at the highest concentration tested. (B) Dose–response curve for the positive control trichostatin A (TSA), yielding an $IC_{50}$ of 3.3 nM, confirming assay performance.

*Compound **B** (2E)-4-(3-phenoxyphenyl)but-2-enoate*

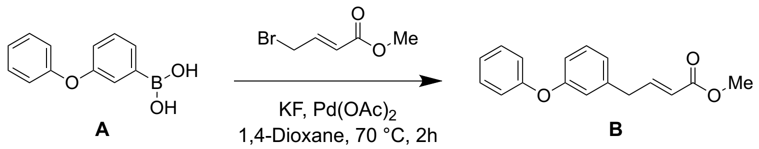


To a stirred mixture of 3-phenoxyphenylboronic acid **A** (20 g, 93.4 mmol) and methyl (2E)-4-bromobut-2-enoate (16.73 g, 93.4 mmol) in 1,4-dioxane (200 mL) were added KF (10.86 g, 186.8 mmol) and $Pd(OAc)_2$ (2.10 g, 9.3mmol) in portions at 25°C under argon atmosphere. The resulting mixture was stirred at 70°C for additional 2h. The reaction was quenched with ice water at 0°C. The resulting mixture was extracted with EtOAc (3 x 30mL). The combined organic layers were washed with brine (2x200 mL), dried over anhydrous $Na_2SO_4$. After filtration, the filtrate was concentrated under reduced pressure. The residue was purified by silica gel column chromatography, eluted with PE / EA (5:1) to afford **B** methyl (2E)-4-(3-phenoxyphenyl)but-2-enoate (13 g, 51.85% yield, 90% purity) as a colorless oil.

$^1$H NMR (400 MHz, DMSO) δ 7.44 – 7.30 (m, 4H), 7.15 – 7.10 (m, 1H), 7.01 (dddd, $J$ = 6.9, 3.4, 2.3, 1.2 Hz, 4H), 6.86 (d, $J$ = 8.1 Hz, 1H), 5.89 (d, $J$ = 15.5 Hz, 1H), 3.64 (d, $J$ = 1.1 Hz, 3H), 3.57 – 3.53 (m, 2H).

*Compound **C** (3S,4S)-1-benzyl-4-[(3-phenoxyphenyl)methyl]pyrrolidine-3-carboxylate*

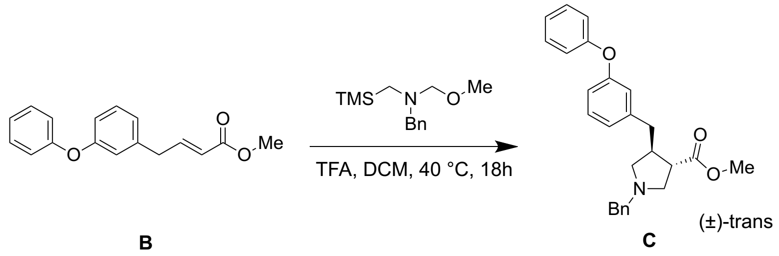


To a stirred mixture of **B** methyl (2E)-4-(3-phenoxyphenyl)but-2-enoate (13 g, 48.451 mmol, 1 equiv) and benzyl(methoxymethyl)[(trimethylsilyl)methyl]amine (20.71 g, 87.2 mmol) in DCM (130 mL) was added TFA (1.66 g, 14.5 mmol) in portions at 0° C under argon atmosphere. The resulting mixture was stirred at 40° C for additional 18h. Desired product could be detected by LCMS. The reaction was quenched with ice water at 0° C. The resulting mixture was extracted with EtOAc (2 x200 mL). The combined organic layers were washed with brine (2x20 mL), dried over anhydrous $Na_2SO_4$. After filtration, the filtrate was concentrated under reduced pressure. The residue was purified by silica gel column chromatography, eluted with PE / EA (5:1) to afford **C** methyl (3S,4S)-1-benzyl-4-[(3-phenoxyphenyl)methyl]pyrrolidine-3-carboxylate (10 g, 51.40%yield, 90%purity) as a white solid.

LCMS-C:(ES,m/z):402.2 $[M+H]^+$

$^1$H NMR (400 MHz, DMSO) δ 7.41 – 7.33 (m, 2H), 7.34 – 7.20 (m, 7H), 7.12 (tt, *J* = 7.3, 1.1 Hz, 1H), 7.00 – 6.92 (m, 2H), 6.85 – 6.78 (m, 2H), 3.62– 3.54 (m, 2H), 3.51 (s, 3H), 2.83 – 2.54 (m, 8H).

*Compound **D** methyl (3S,4S)-4-[(3-phenoxyphenyl)methyl]pyrrolidine-3-carboxylate*

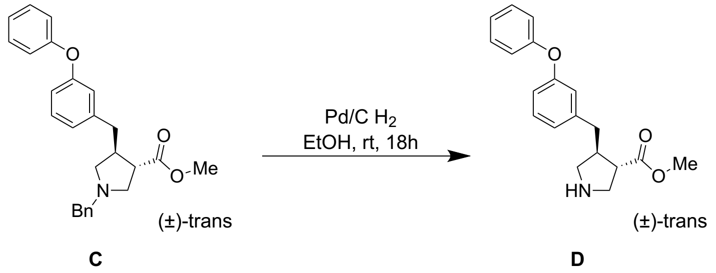


To a stirred mixture of **C** methyl (3S,4S)-1-benzyl-4-[(3-phenoxyphenyl)methyl]pyrrolidine-3-carboxylate (10 g, 24.9 mmol) in EtOH (100 mL) was added Pd/C (10 g, 93.9 mmol) in portions at 0° C under hydrogen atmosphere. The resulting mixture **D** methyl (3S,4S)-4-[(3-phenoxyphenyl)methyl]pyrrolidine-3-carboxylate was stirred at 25° C for additional overnight. Desired product could be detected by LCMS. The resulting mixture was filtered, the filter cake was washed with EtOH (100 mL) (2x200 mL). The filtrate was concentrated under reduced pressure. The resulting mixture was used in the next step directly without further purification.

LCMS-D:(ES,m/z):312.2 $[M+H]^+$

$^1$H NMR (400 MHz, DMSO) δ 7.39 (t, *J* = 7.8 Hz, 2H), 7.30 (t, *J* = 7.8 Hz, 1H), 7.13 (t, *J* = 7.4 Hz, 1H), 6.98 (t, *J* = 7.0 Hz, 3H), 6.92 – 6.79 (m, 2H), 3.51 (s, 3H), 3.28 – 3.08 (m, 2H), 2.94 (dt, *J* = 7.2, 4.8 Hz, 2H), 2.72 – 2.53 (m, 4H).

*Compound **E** 1-tert-butyl 3-methyl (3S,4S)-4-[(3-phenoxyphenyl)methyl]pyrrolidine-1,3-dicarboxylate*

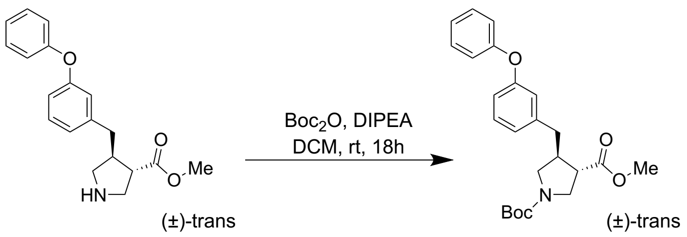


**D** **E**

To a stirred mixture of **D** methyl (3S,4S)-4-[(3-phenoxyphenyl)methyl]pyrrolidine-3-carboxylate (6 g, 19.269 mmol, 1 equiv) and DIEA (4.98 g, 38.5 mmol) in DCM (60 mL) was added Boc$_2$O (4.21 g, 19.2 mmol) in portions at 0° C under air atmosphere. The resulting mixture was stirred at 25° C for additional overnight. Desired product could be detected by LCMS. The reaction was quenched with ice water at 0° C. The resulting mixture was extracted with EtOAc (2 x 100mL). The combined organic layers were washed with hexane (2x100 mL), dried over anhydrous Na2SO4. After filtration, the filtrate was concentrated under reduced pressure. The residue was purified by silica gel column chromatography, eluted with PE / EA (4:1) to afford **E** 1-tert-butyl 3-methyl (3S,4S)-4-[(3-phenoxyphenyl)methyl]pyrrolidine-1,3-dicarboxylate (4.2 g, 52.97%yield, 90%purity) as a colorless oil.

LCMS- E:(ES,m/z):412.2 $[M+H]^+$

$^1$H NMR (400 MHz, DMSO) δ 7.42 – 7.35 (m, 2H), 7.31 (t, *J* = 7.8 Hz, 1H), 7.16 – 7.10 (m, 1H), 7.03 – 6.95 (m, 3H), 6.90 – 6.81 (m, 2H), 3.65 – 3.57 (m, 1H), 3.54 (d, *J* = 2.0 Hz, 3H), 3.35 (d, *J* = 7.1 Hz, 2H), 2.97 (q, *J* = 8.7 Hz, 1H), 2.92 – 2.73 (m, 2H), 2.63 (dt, *J* = 19.4, 7.6 Hz, 2H), 1.37 (d, *J* = 2.7 Hz, 9H).

*Compound* ***F*** *(3S,4S)-1-(tert-butoxycarbonyl)-4-[(3-phenoxyphenyl)methyl]pyrrolidine-3-carboxylic acid*

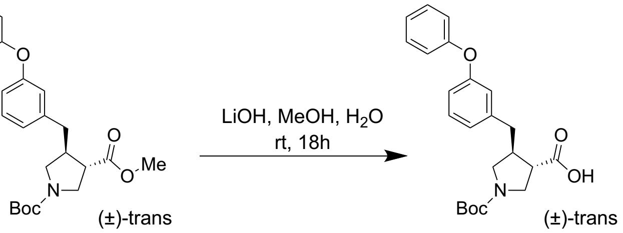


**E** **F**

To a stirred mixture of **E** 1-tert-butyl 3-methyl (3S,4S)-4-[(3-phenoxyphenyl)methyl]pyrrolidine-1,3-dicarboxylate (5.2 g, 12.6 mmol) and MeOH (50 mL) in $H_2O$ (12 mL) was added LiOH (1.51 g, 63.1 mmol) in portions at 25° C under air atmosphere. The resulting mixture was stirred at 25° C for additional overnight. Desired product **F** (3S,4S)-1-(tert-butoxycarbonyl)-4-[(3-phenoxyphenyl)methyl]pyrrolidine-3-carboxylic acid could be detected by LCMS. The mixture to pH 4 with conc. HCl. The resulting mixture was extracted with EtOAc (2 x 200mL). The combined organic layers were washed with brine (1x100 mL), dried over anhydrous $Na_2SO_4$. After filtration, the filtrate was concentrated under reduced pressure. The crude product/ resulting mixture was used in the next step directly without further purification.

LCMS- F:(ES,m/z):398.2 $[M+H]^+$

$^1H$ NMR (400 MHz, DMSO) δ 12.51 (s, 1H), 7.42 – 7.35 (m, 2H), 7.31 (t, *J* = 7.8 Hz, 1H), 7.13 (td, *J* = 7.3, 1.1 Hz, 1H), 6.99 (ddd, *J* = 7.6, 3.0, 1.7 Hz, 3H), 6.93 – 6.80 (m, 2H), 3.53 (dtt, *J* = 14.5, 10.4, 4.9 Hz, 2H), 3.38 (d, *J* = 8.3 Hz, 1H), 3.27 (d, *J* = 6.8 Hz, 1H), 2.97 (q, *J* = 8.1 Hz, 1H), 2.89 – 2.78 (m, 1H), 2.74 (q, *J* = 6.9 Hz, 1H), 2.61 (t, *J* = 8.9 Hz, 2H), 1.45 – 1.32 (m, 13H).

*Compound **G** tert-butyl (3R,4R)-3-[(4S)-4-(2-methylidenebut-3-en-1-yl)-2-oxo-1,3-oxazolidine-3-carbonyl]-4-[(3-phenoxyphenyl)methyl]pyrrolidine-1-carboxylate*

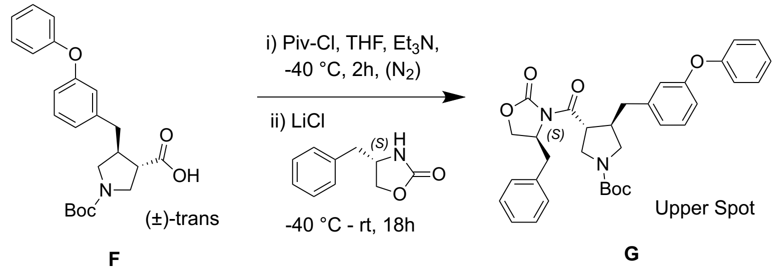


To a stirred mixture of **F** (3S,4S)-1-(tert-butoxycarbonyl)-4-[(3-phenoxyphenyl)methyl]pyrrolidine-3-carboxylic acid (4.6 g, 11.5 mmol) and $Et_3N$ (3.51 g, 34.719 mmol) in THF (50 mL) was added 2,2-dimethylpropanoyl chloride (1.54 g, 12.7 mmol) in portions at -40° C under argon atmosphere. The resulting mixture was stirred at -40° C for additional 2h. To the above mixture was added LiCl (20.60 g, 486.066 mmol) and (4S)-4-benzyl-1,3-oxazolidin-2-one (2.05 g, 11.5 mmol) in portions over 1min at -40° C. The resulting mixture was stirred at 25° C for additional overnight. Desired product could be detected by

LCMS. The reaction was quenched with sat. $NH_4Cl$ (aq.) at 0° C. The resulting mixture was extracted with EtOAc (2 x 100mL). The combined organic layers were washed with brine (2x200 mL), dried over anhydrous Na2SO4. After filtration, the filtrate was concentrated under reduced pressure. The residue was purified by silica gel column chromatography, eluted with PE / EA (4:1) to afford **G** tert-butyl (3R,4R)-3-[(4S)-4-(2-methylidenebut-3-en-1-yl)-2-oxo-1,3-oxazolidine-3-carbonyl]-4-[(3-phenoxyphenyl)methyl]pyrrolidine-1-carboxylate (1.5 g, 24.33%yield, 90%purity) as a white solid.

LCMS- G:(ES,m/z):557.3 $[M+H]^+$

$^1$H NMR (400 MHz, DMSO) δ 7.42 – 7.35 (m, 2H), 7.34 – 7.23 (m, 4H), 7.14 (q, *J* = 6.7 Hz, 3H), 7.00 (d, *J* = 8.1 Hz, 2H), 6.95 (d, *J* = 7.6 Hz, 1H), 6.84 (d, *J* = 3.4 Hz, 2H), 4.50 (s, 1H), 4.30 (t, *J* = 8.5 Hz, 1H), 4.23 (dd, *J* = 8.8, 2.9 Hz, 1H), 3.82 (dd, *J* = 24.3, 7.1 Hz, 1H), 3.71 – 3.58 (m, 1H), 3.37 (dd, *J* = 10.3, 6.9 Hz, 2H), 3.09 – 3.01 (m, 1H), 2.95 (d, *J* = 5.8 Hz, 2H), 2.82 (dd, *J* = 13.3, 6.6 Hz, 1H), 2.71 (dd, *J* = 13.4, 6.3 Hz, 1H), 2.62 (dd, *J* = 13.5, 8.8 Hz, 1H), 1.40 (d, *J* = 9.5 Hz, 9H).

*Compound **H** (3R,4R)-1-(tert-butoxycarbonyl)-4-[(3-phenoxyphenyl)methyl]pyrrolidine-3-carboxylic acid*

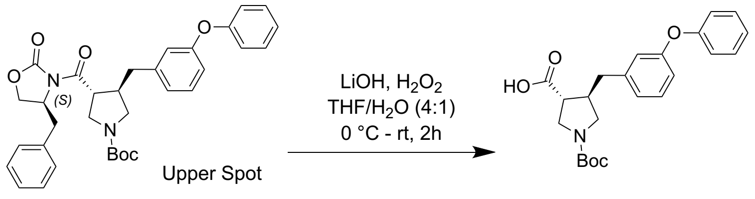


**G** **H**

A solution of **G** tert-butyl (3R,4R)-3-[(4S)-4-benzyl-2-oxo-1,3-oxazolidine-3-carbonyl]-4-[(3-phenoxyphenyl)methyl]pyrrolidine-1-carboxylate (250 mg, 0.4 mmol) in THF (2 mL) was treated with $H_2O$ (0.5 mL, 0.4 mmol) at 0° C for 1min under nitrogen atmosphere followed by the addition of LiOH (21.51 mg, 0.898 mmol) in portions at 0° C. The resulting mixture was stirred at room temperature for additional 3h. Desired product could be detected by LCMS. The residue was acidified to pH 6 with conc. HCl. The resulting mixture was extracted with EtOAc (2 x 10mL). The combined organic layers were washed with hexane (2x10 mL), dried over anhydrous $Na_2SO_4$. After filtration, the filtrate was concentrated under reduced pressure. The crude product was purified by Prep-HPLC with the following conditions (Column: Xselect CSH

Prep C18 30*150mm 5 μ m; Mobile Phase A: Water(0.1% FA), Mobile Phase B: ACN; Flow rate: 60 mL/min mL/min; Gradient (B%): 54% B to 74% B in 8 min; Wave Length: 254nm/220nm nm; RT1(min): 8.1) to afford **H** (3R,4R)-1-(tert-butoxycarbonyl)-4-[(3-phenoxyphenyl)methyl]pyrrolidine-3-carboxylic acid (105 mg, 58.82%yield, 99.5%purity) as a colorless oil.

LCMS- H :(ES,m/z):398.1$[M+1]^+$

$^1$H NMR (400 MHz, DMSO) δ 12.56 (s, 1H), 7.38 (t, $J$ = 7.7 Hz, 2H), 7.31 (t, $J$ = 7.8 Hz, 1H), 7.13 (t, $J$ = 7.4 Hz, 1H), 6.99 (dd, $J$ = 8.2, 3.6 Hz, 3H), 6.91 – 6.75 (m, 2H), 3.55 (t, $J$ = 9.7 Hz, 1H), 3.38 (s, 2H), 2.97 (q, $J$ = 8.1 Hz, 1H), 2.84 (s, 1H), 2.73 (q, $J$ = 6.9 Hz, 1H), 2.60 (q, $J$ = 7.1 Hz, 2H), 1.37 (d, $J$ = 4.7 Hz, 9H).

*Compound **I** tert-butyl (3R,4R)-3-((2-((2-(dimethylamino)ethyl)disulfaneyl)ethyl)carbamoyl)-4-(3-phenoxybenzyl)pyrrolidine-1-carboxylate*

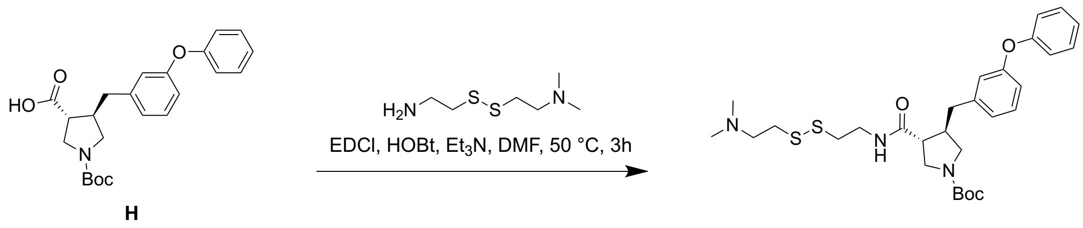


To a stirred mixture of **H** (3R,4R)-1-(tert-butoxycarbonyl)-4-[(3-phenoxyphenyl)methyl]pyrrolidine-3-carboxylic acid (450 mg, 1.13 mmol) and (2-[(2-aminoethyl)disulfanyl]ethyldimethylamine (203.4 mg, 1.13 mmol) in DMF (1.0mL) were added $Et_3N$ (460 mg, 4.51 mmol) and HOBt (184 mg, 1.36 mmol,) , EDCI (211 mg, 1.31 mmol,) in portions at 0° C under argon atmosphere. The resulting mixture was stirred at 50° C for additional 3h. Desired product could be detected by LCMS. The residue was purified by reversed-phase flash chromatography with the following conditions: column, C18 silica gel; mobile phase, MeCN in Water (10 mmol/L $NH_4HCO_3$), 10% to 50% gradient in 10 min; detector, UV 254 nm. This resulted in **I** tert-butyl (3R,4R)-3-((2-((2-(dimethylamino)ethyl)disulfaneyl)ethyl)carbamoyl)-4-(3-phenoxybenzyl)pyrrolidine-1-carboxylate (350 mg, 55.2%yield) as a yellow solid.

LCMS- I :(ES,m/z):560.2[M+1]+

[1]H NMR (400 MHz, DMSO) δ 8.22 (t, $J$ = 5.7 Hz, 1H), 7.38 (t, $J$ = 7.7 Hz, 2H), 7.29 (t, $J$ = 7.9 Hz, 1H), 7.13 (tt, $J$ = 7.3, 1.1 Hz, 1H), 7.04 – 6.92 (m, 3H), 6.83 (d, $J$ = 11.9 Hz, 2H), 3.50 (dd, $J$ = 10.6, 7.9 Hz, 1H), 3.37 – 3.18 (m, 5H), 2.91 (dd, $J$ = 12.2, 7.6 Hz, 1H), 2.84 (dd, $J$ = 8.1, 6.2 Hz, 2H), 2.76 (t, $J$ = 6.7 Hz, 3H), 2.61 – 2.53 (m, 3H), 2.20 (s, 6H).

*Compound **J** (3R,4R)-N-(2-([2-(dimethylamino)ethyl]disulfanylethyl)-4-[(3-phenoxyphenyl)methyl]pyrrolidine-3-carboxamide*

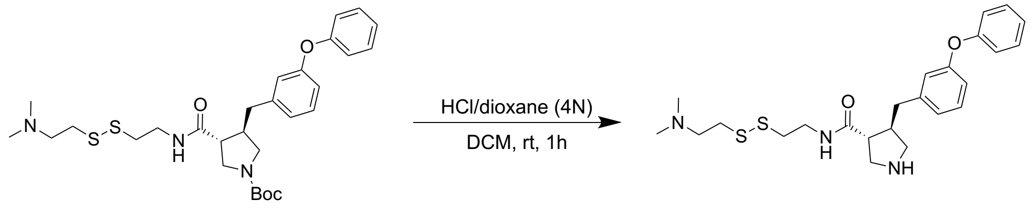


**I** **J**

To a stirred mixture of **I** tert-butyl (3R,4R)-3-[(2-([2-(dimethylamino)ethyl]disulfanylethyl)carbamoyl]-4-[(3-phenoxyphenyl)methyl]pyrrolidine-1-carboxylate (390 mg, 0.6 mmol) in DCM (4 mL) was added HCl in 1,4-dioxane (4.0 M) (2 mg, 0.05 mmol) in portions at 0° C under air atmosphere. The resulting mixture was stirred at 0° C for additional 2h.The resulting mixture **J** (3R,4R)-N-(2-([2-(dimethylamino)ethyl]disulfanylethyl)-4-[(3-phenoxyphenyl)methyl]pyrrolidine-3-carboxamide was concentrated under reduced pressure. The resulting mixture was used in the next step directly without further purification.

LCMS- J:(ES,m/z):460.2[M+1]+

***Compound 1*** *(3R,4R)-1-(3,5-dichloro-4-hydroxybenzenesulfonyl)-N-(2-([2-(dimethylamino)ethyl]disulfanylethyl)-4-[(3-phenoxyphenyl)methyl]pyrrolidine-3-carboxamide*

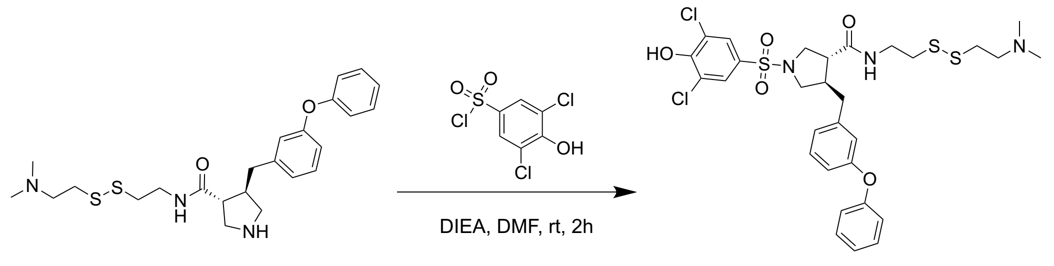


**J** **Compound 1**

To a stirred mixture of **J** (3R,4R)-N-(2-([2-(dimethylamino)ethyl]disulfanylethyl)-4-[(3-phenoxyphenyl)methyl]pyrrolidine-3-carboxamide (150 mg, 0.3 mmol) and DIEA (84.35 mg, 0.6 mmol) in DMF (2 mL) were added 3,5-dichloro-4-hydroxybenzenesulfonyl chloride (85.33 mg, 0.3 mmol) in portions at 0° C under air atmosphere. The resulting mixture was stirred at 0 ° C for additional 2h. Desired product could be detected by LCMS. The resulting mixture was concentrated under reduced pressure. The crude product was purified by Prep-HPLC with the following conditions (Column: XBridge Shield RP18 OBD Column 30*150 mm, 5 μ m; Mobile Phase A: Water(10mmol/L NH4HCO3), Mobile Phase B: ACN; Flow rate: 60 mL/min; Gradient (B%): 32% B to 45% B in 11 min; Wave Length: 254nm/220 nm; RT1(min): 10.77) to afford **Compound 1** (3R,4R)-1-(3,5-dichloro-4-hydroxybenzenesulfonyl)-N-(2-([2-(dimethylamino)ethyl]disulfanylethyl)-4-[(3-phenoxyphenyl)methyl]pyrrolidine-3-carboxamide (18 mg, 8.06%yield, 97.9%purity) as a white solid.

HRMS (ESI-TOF) m/z: calcd for $C_{30}H_{36}Cl_2N_3O_5S_3$ $[M + H]^+$, 684.1189; found, 684.1201.

$[\alpha]_D^{20}$ = +13.3 (c 0.30, $CH_2Cl_2$/MeOH, 9:1).

$^1$H NMR (400 MHz, $CDCl_3$) δ 7.60 (s, 1H), 7.38 – 7.29 (m, 2H), 7.24 – 7.16 (m, 1H), 7.09 (tt, *J* = 7.4, 1.1 Hz, 1H), 7.00 – 6.91 (m, 3H), 6.85 – 6.77 (m, 3H), 3.51 (t, *J* = 9.1 Hz, 1H), 3.35 (qd, *J* = 14.4, 6.9 Hz, 4H), 3.27 – 3.18 (m, 5H), 3.19 – 2.83 (m, 3H), 2.77 – 2.64 (m, 7H), 2.62 – 2.37 (m, 3H).

$^{13}$C NMR (101 MHz, $CDCl_3$) δ 171.67, 161.37, 157.41, 157.05, 141.18, 129.91, 129.84, 127.88, 124.75, 123.71, 123.38, 120.32, 119.21, 118.91, 116.83, 77.25, 57.40, 52.78, 51.06, 49.31, 44.83, 43.78, 42.97, 38.50, 38.40, 37.51, 32.05.

***Compound 2*** *(3R,4R)-1-(3,5-dichloro-4-hydroxybenzoyl)-N-(2-([2-(dimethylamino)ethyl]disulfanylethyl)-4-[(3-phenoxyphenyl)methyl]pyrrolidine-3-carboxamide*

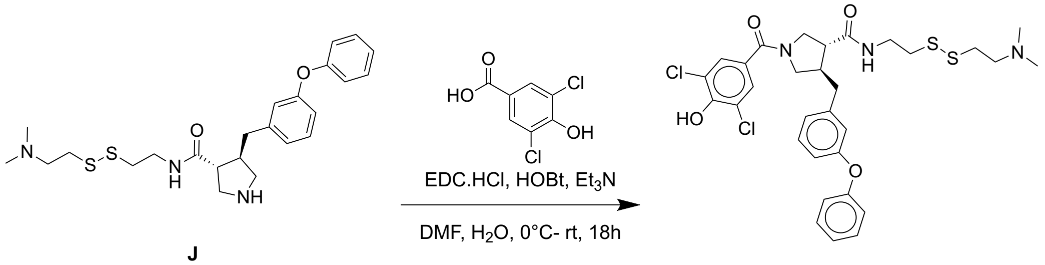


**J**

**Compound 2**

To a stirred mixture of **J** (3R,4R)-N-(2-([2-(dimethylamino)ethyl]disulfanylethyl)-4-[(3-phenoxyphenyl)methyl]pyrrolidine-3-carboxamide (110 mg, 0.2 mmol) and 3,5-dichloro-4-hydroxybenzoic acid (41.2 mg, 0.2 mmol) in DMF (2 mL) were added $Et_3N$ (48.43 mg, 0.4 mmol) and EDCI (37.15 mg, 0.2 mmol) in portions at 0° C under nitrogen atmosphere. To the above mixture was added HOBt (27 mg, 0.2 mmol) in portions over 1min at 0° C. The resulting mixture was stirred at 0° C for additional 2h. Desired product could be detected by LCMS. The crude product was purified by Prep-HPLC with the following conditions (Column: XBridge Prep OBD C18 Column 30*150 mm, 5 μ m; Mobile Phase A: Water(10mmol/L NH4HCO3), Mobile Phase B: ACN; Flow rate: 60 mL/min; Gradient (B%): 30% to 44% B in 10 min; RT1(min): 8.72) to afford **Compound 2** (3R,4R)-1-(3,5-dichloro-4-hydroxybenzoyl)-N-(2-([2-(dimethylamino)ethyl]disulfanylethyl)-4-[(3-phenoxyphenyl)methyl]pyrrolidine-3-carboxamide (21 mg, 13.53%yield, 96.5%purity) as a white solid.

HRMS (ESI-TOF) m/z: calcd for $C_{31}H_{36}Cl_2N_3O_4S_2$ $[M + H]^+$, 648.1519**;** found, 648.1558**.**

$[\alpha]_D^{20}$ = +75.38 (c 0.325, $CH_2Cl_2$/MeOH, 9:1).

NMR-Compound 2: 1H NMR (400 MHz, DMSO) δ 8.26 (t, *J* = 5.7 Hz, 1H), 7.36 (dt, *J* = 8.3, 7.0 Hz, 2H), 7.27 (t, *J* = 7.9 Hz, 1H), 7.21 (s, 1H), 7.15 – 7.07 (m, 1H), 7.04 – 6.90 (m, 3H), 6.89 – 6.75 (m, 2H), 3.69 (s, 1H), 3.60 – 3.46 (m, 2H), 3.28 (td, *J* = 13.8, 6.6 Hz, 4H), 2.87 – 2.71 (m, 5H), 2.65 (q, *J* = 8.1 Hz, 1H), 2.62 – 2.54 (m, 2H), 2.46 (t, *J* = 7.1 Hz, 2H), 2.13 (d, *J* = 10.3 Hz, 6H).

$^{13}C$ NMR (101 MHz, DMSO) δ 167.97, 163.64, 157.13, 157.04, 142.50, 130.48, 130.46, 130.32, 128.32, 124.27, 123.81, 123.79, 122.96, 119.32, 119.00, 116.79, 112.82, 112.72, 58.70, 58.62, 45.38, 45.35, 40.62, 40.41, 38.47, 37.73, 37.55, 36.93, 36.83.

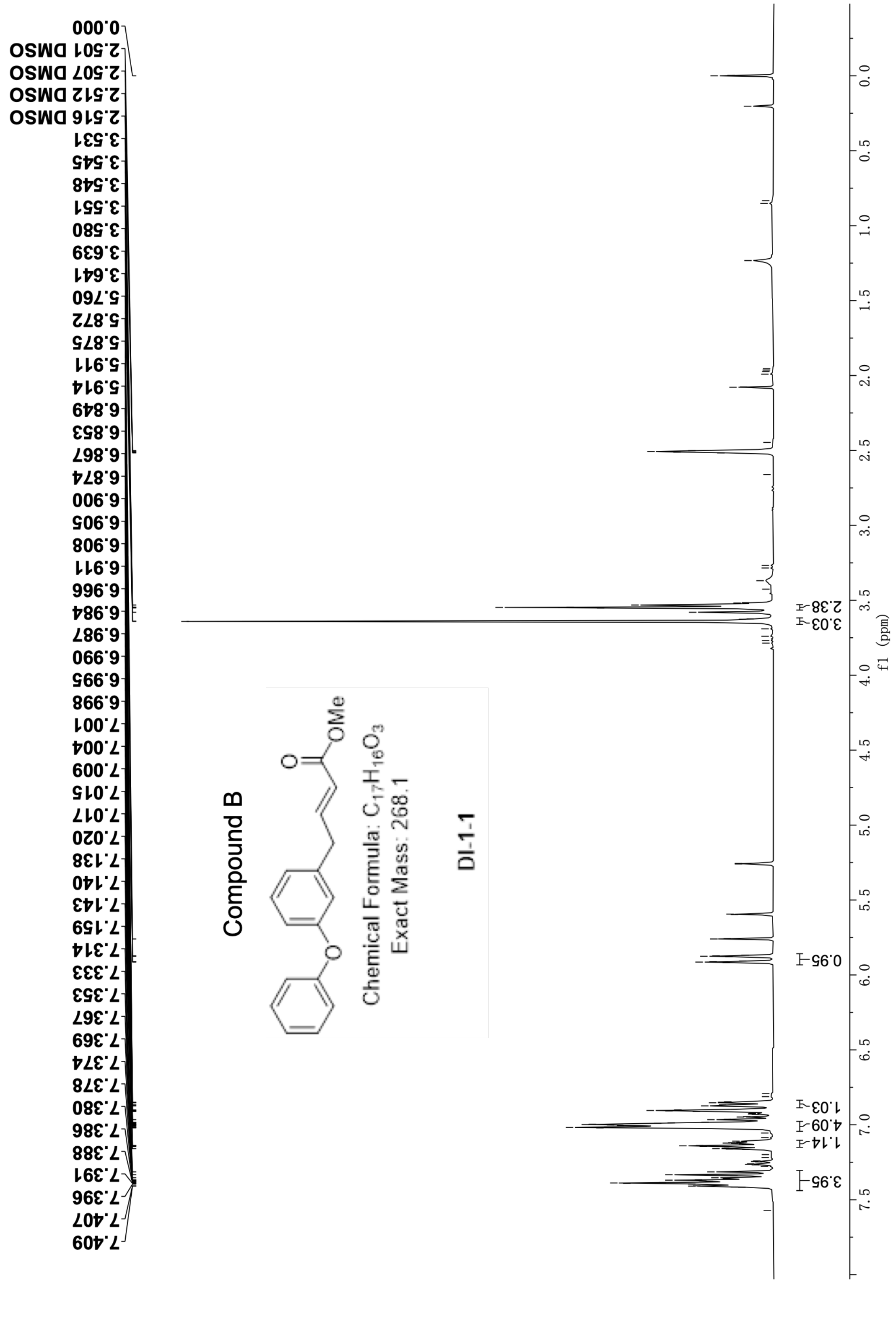


**Figure S.13** H NMR spectrum of Compound B (400 MHz, DMSO-$d_6$).

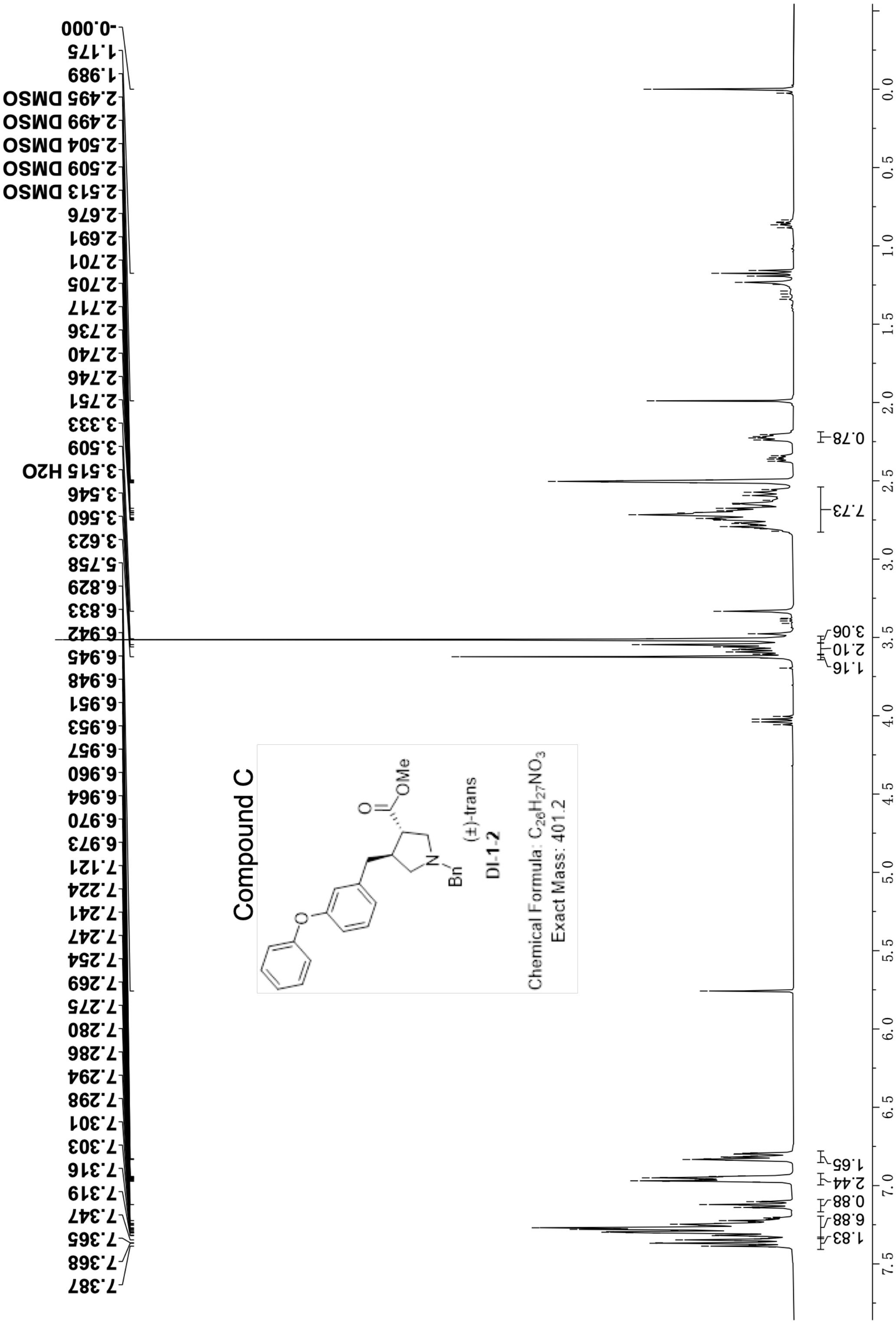


**Figure S.14** H NMR spectrum of Compound C (400 MHz, DMSO-$d_6$).

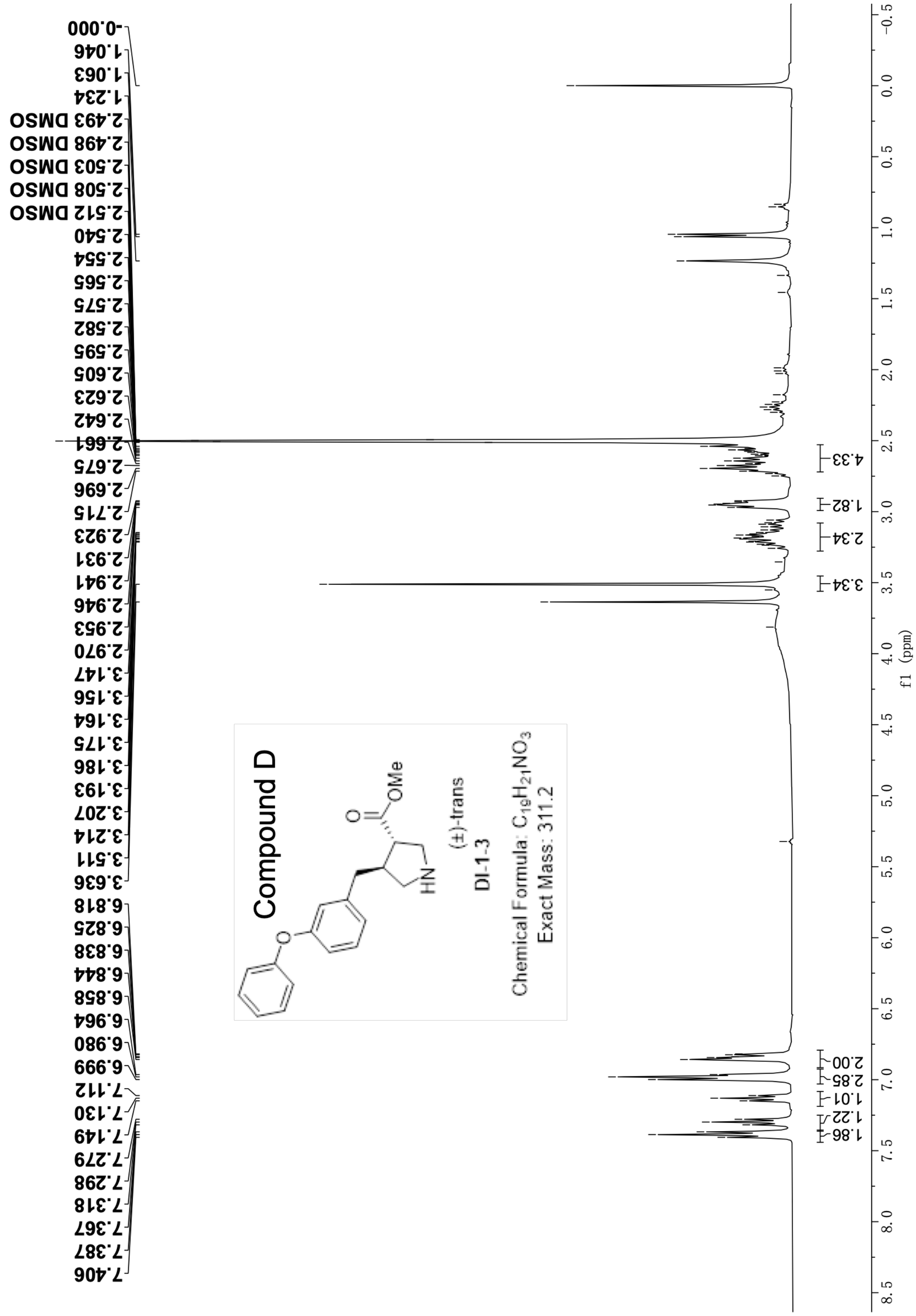


**Figure S.15** H NMR spectrum of Compound D (400 MHz, DMSO-$d_6$).

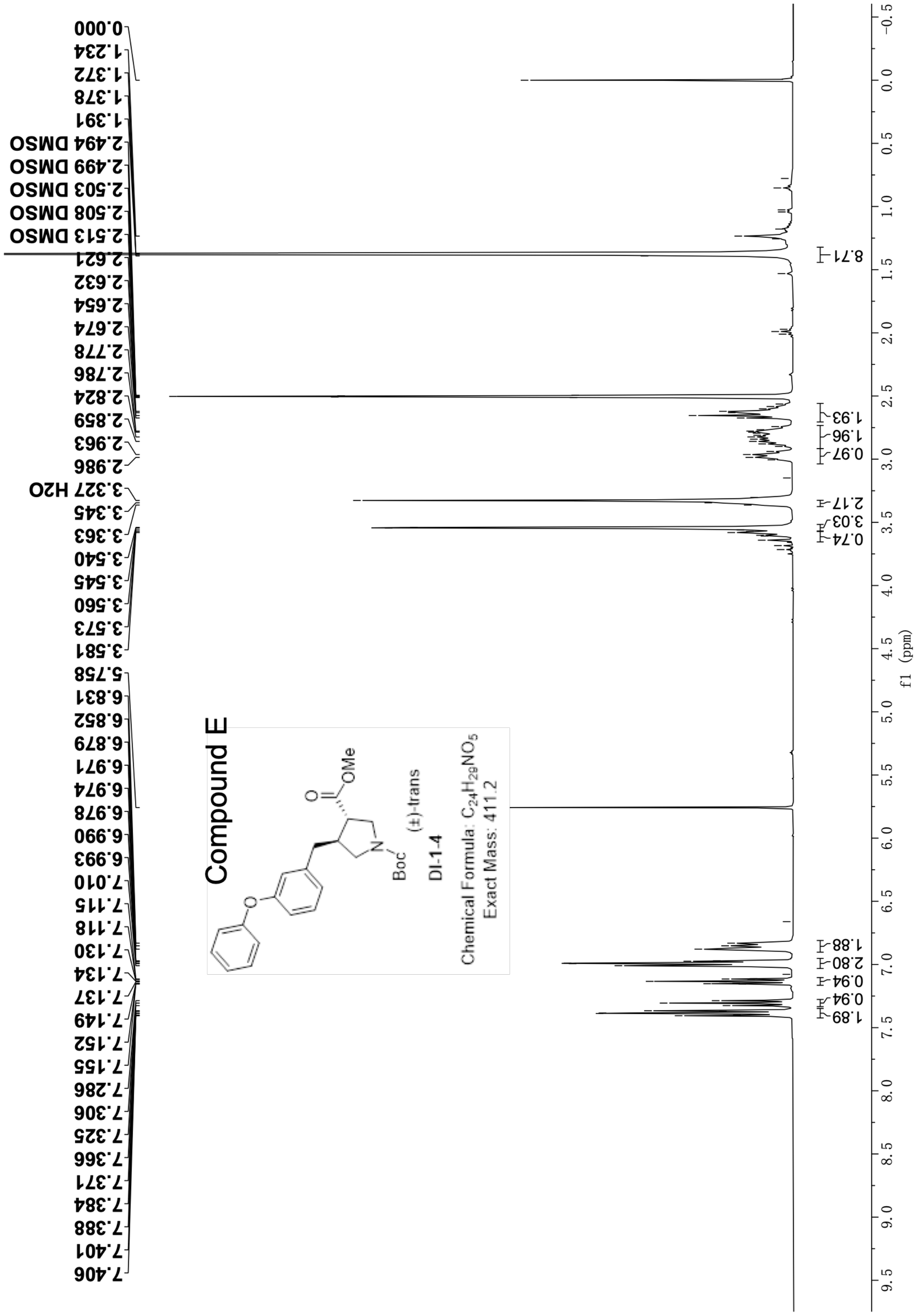


Figure S.16 H NMR spectrum of Compound E (400 MHz, DMSO-$d_6$).

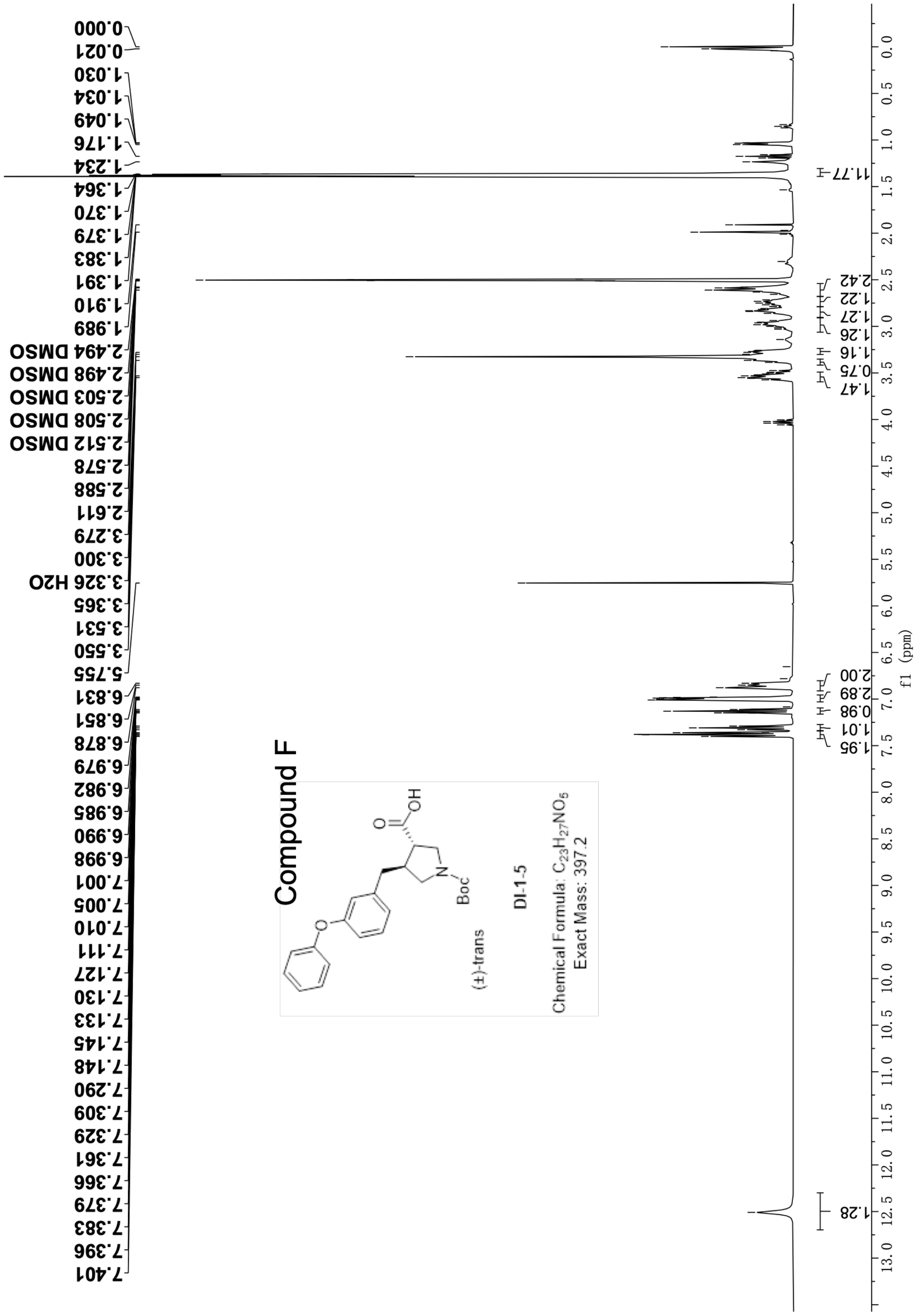


**Figure S.17 H NMR spectrum of Compound F (400 MHz, DMSO-*d*₆).**

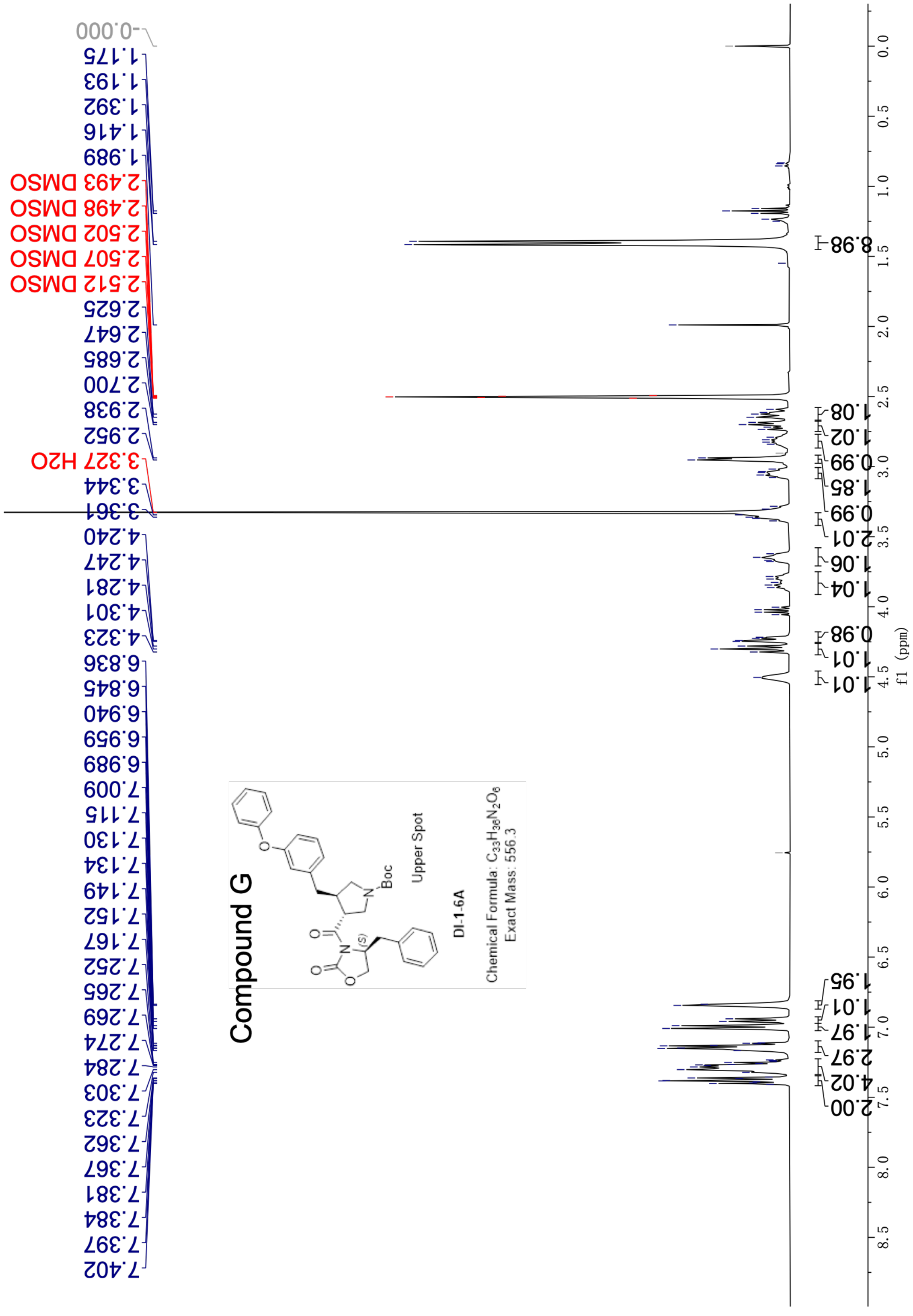


**Figure S.18 H NMR spectrum of Compound G (400 MHz, DMSO-*d*₆).**

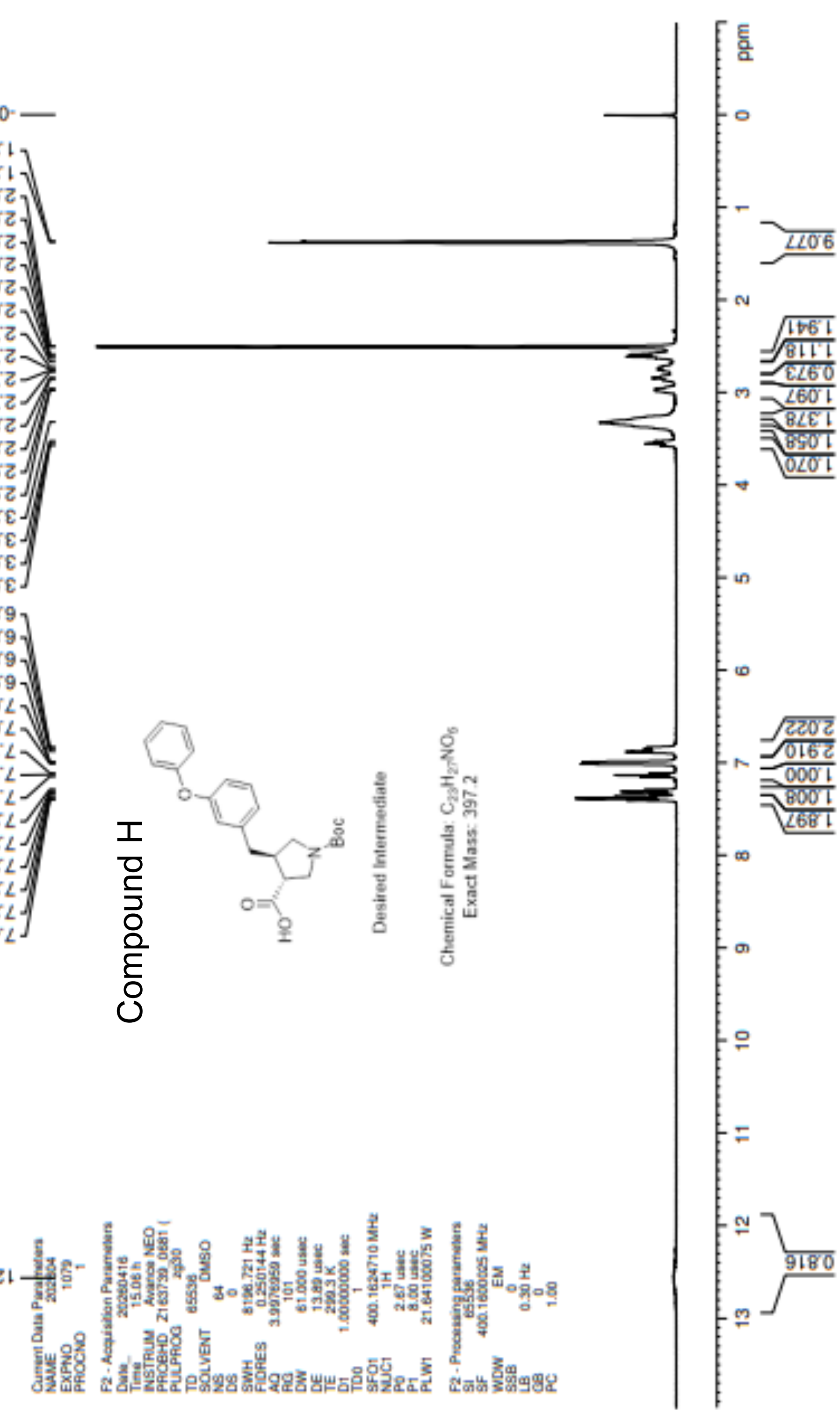


**Figure S.19 H NMR spectrum of Compound H (400 MHz, DMSO-$d_6$).**

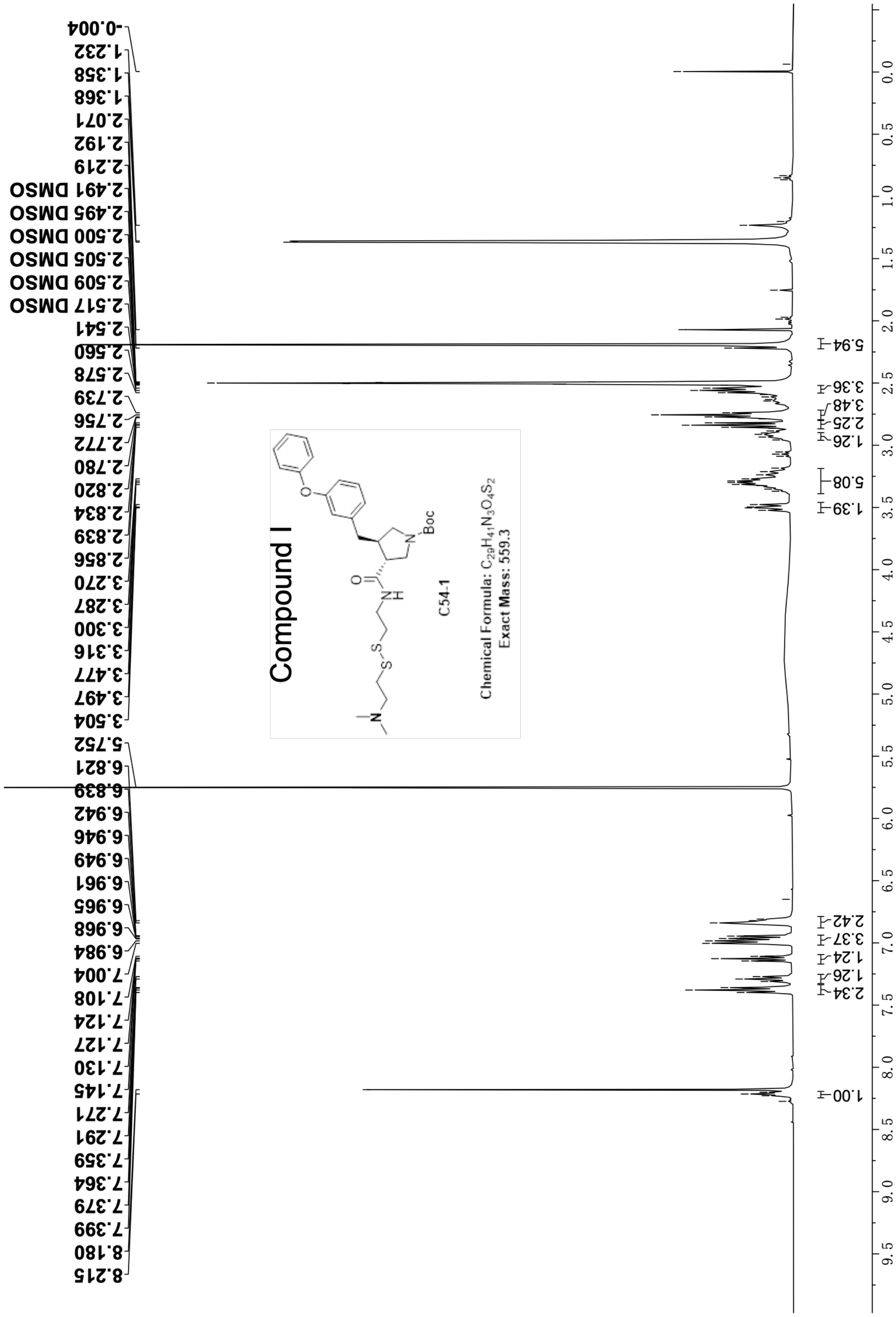


**Figure S.20 H NMR spectrum of Compound I (400 MHz, DMSO-$d_6$).**

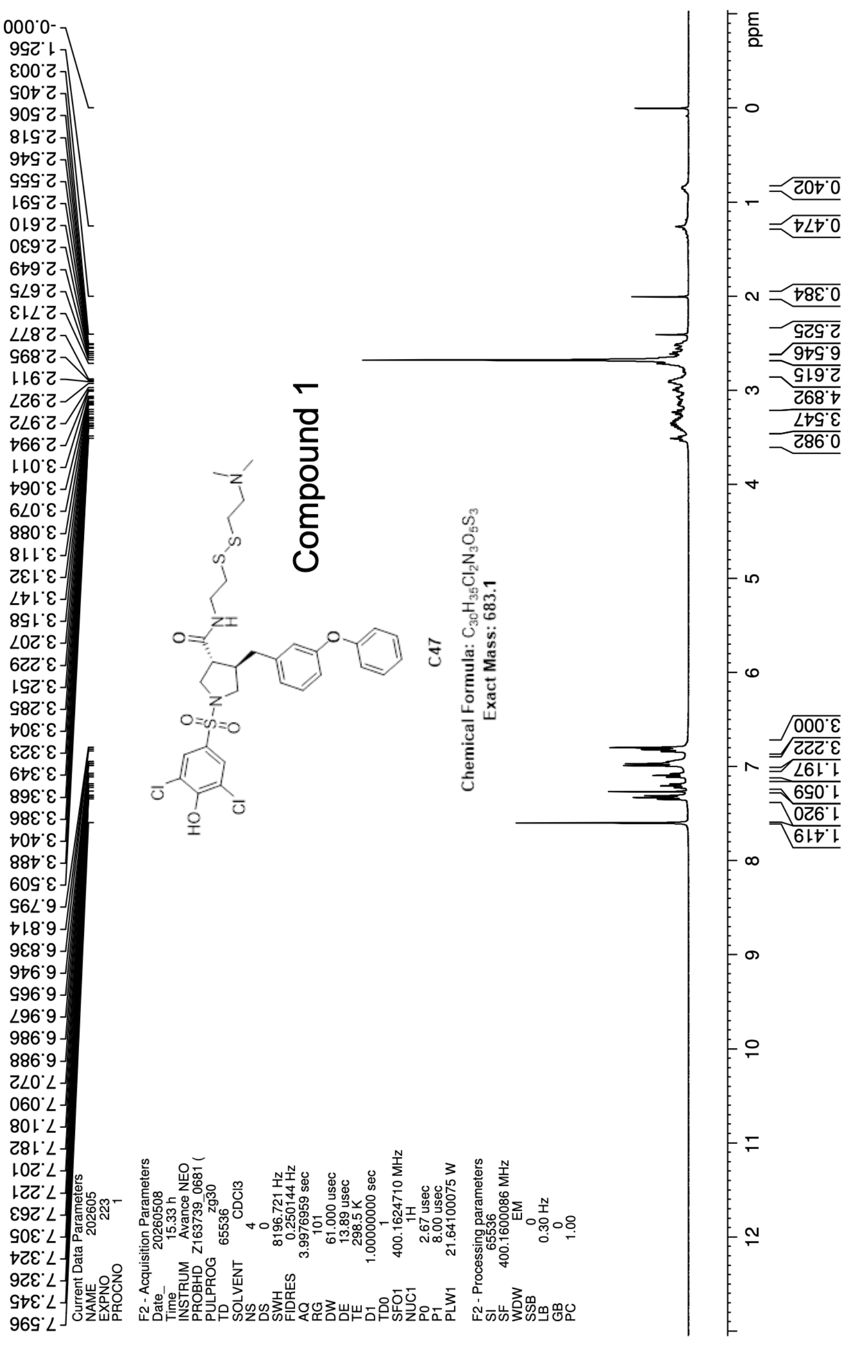


**Figure S.21 H NMR spectrum of Compound 1 (400 MHz, $CDCl_3$).**

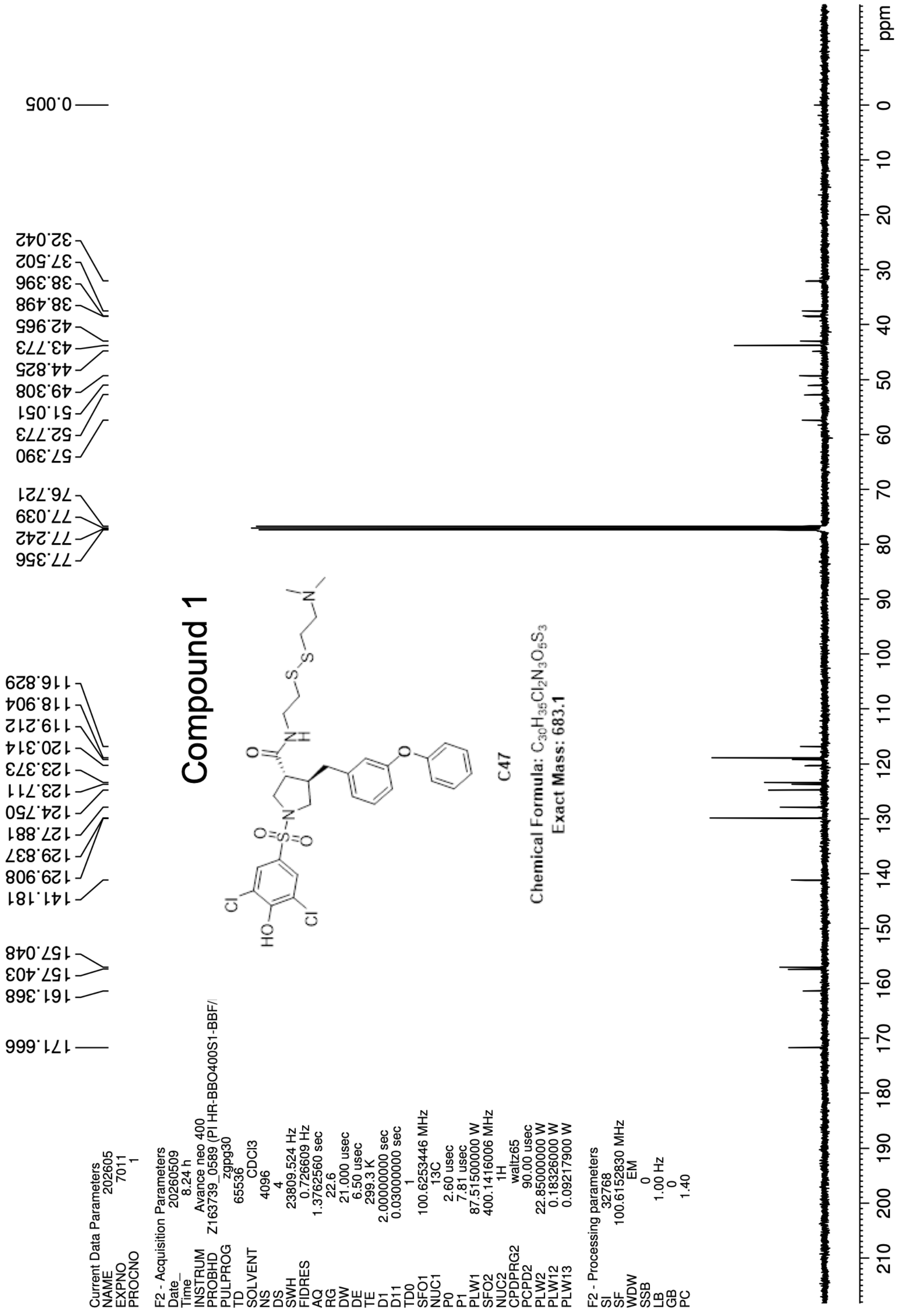


**Figure S.22 C NMR spectrum of Compound 1 (101MHz, CDCl$_3$).**

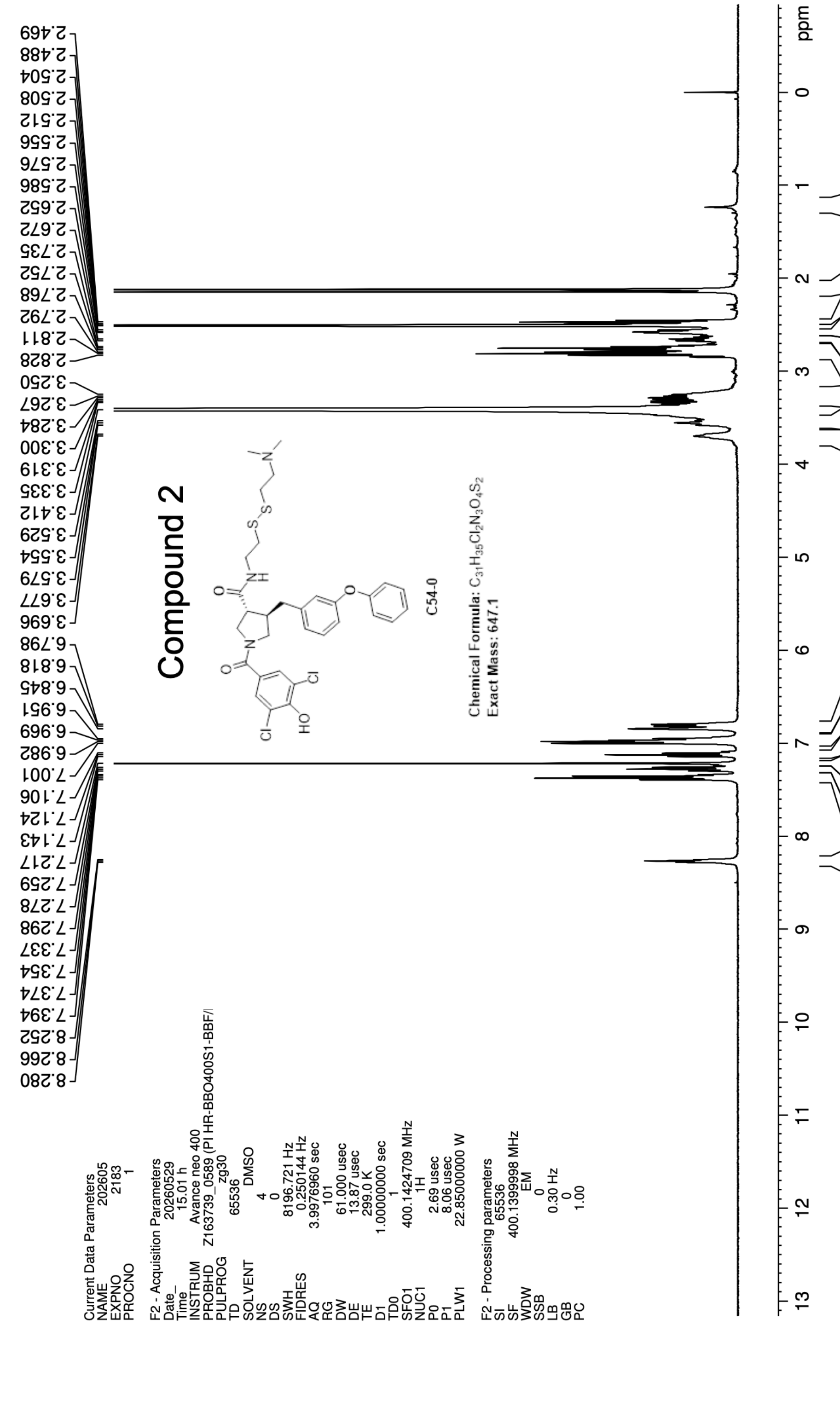


**Figure S.23 H NMR spectrum of Compound 2 (400 MHz, $CDCl_3$).**

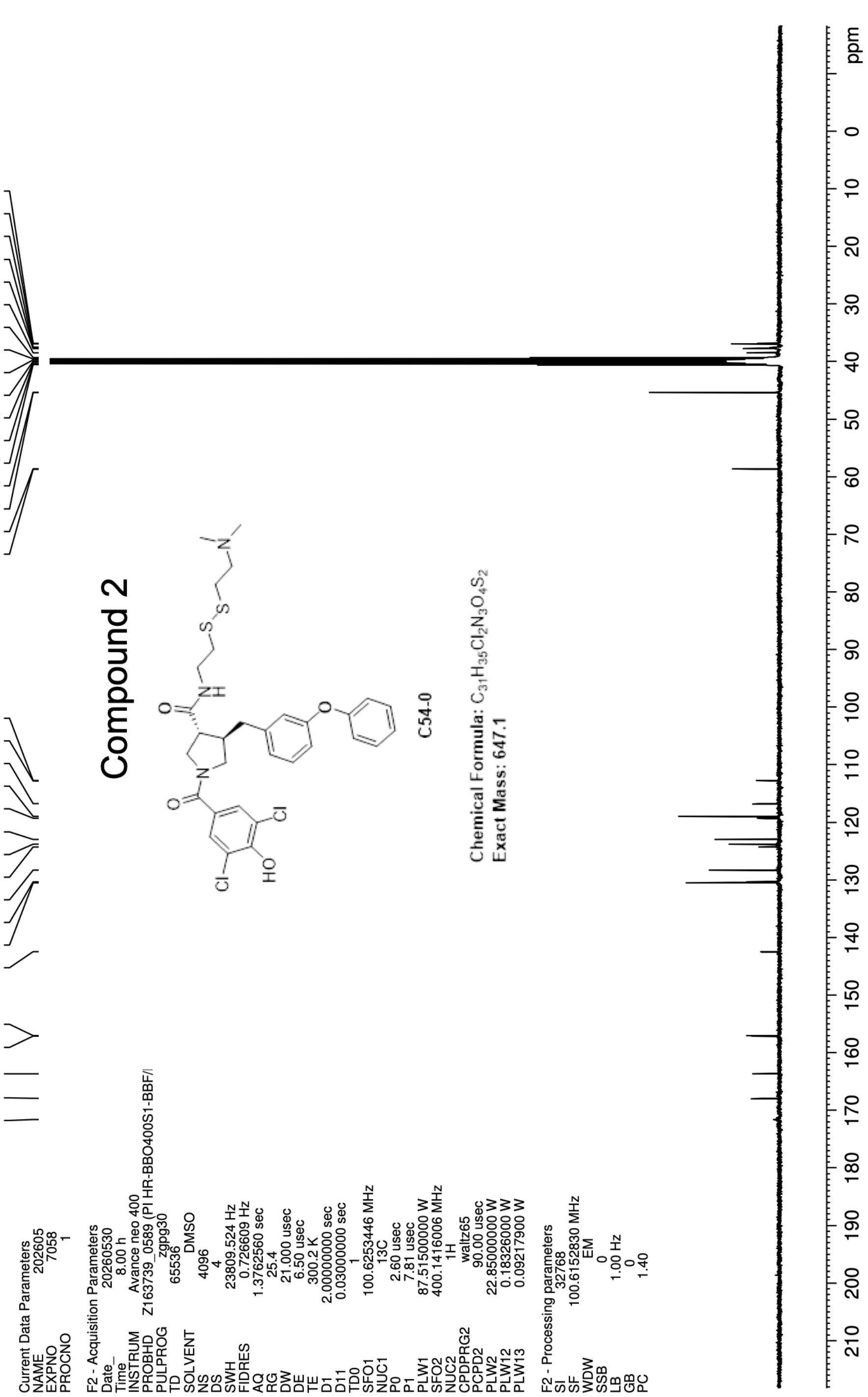


**Figure S.24 C NMR spectrum of Compound 2 (101 MHz, $CDCl_3$).**
